# Diet and chemical defenses of the Sonoran Desert toads

**DOI:** 10.1101/2023.10.06.561297

**Authors:** Marina D. Luccioni, Jules T. Wyman, Edgard O. Espinoza, Lauren A. O’Connell

**Author notes:** These authors contributed equally to this work (Marina D. Luccioni, Jules T. Wyman).

## Abstract

The Sonoran Desert Toad (*Incilius alvarius*) is the only animal known to secrete the psychedelic compound 5-MeO-DMT as a chemical defense, but the source of 5-MeO-DMT in *I. alvarius* remains unknown. Some amphibians endogenously produce chemical defenses while others acquire them from specialized diets. In this study we analyzed toxin gland secretions and diet profiles from wild *I. alvarius* and sympatric anurans from native and urban habitats around Tucson, Arizona to explore possible links between diet and 5-MeO-DMT production. All *I. alvarius* secreted high concentrations of 5-MeO-DMT, whereas other sympatric toads did not. The diet of *I. alvarius* was similar to that of sympatric anurans, indicating that *I. alvarius* does not exhibit relative dietary specialization. Slight dietary differences between *I. alvarius* in native and urbanized habitats were observed. Taken together, these lines of evidence suggest that diet is not directly linked to 5-MeO-DMT production, and support the alternative hypotheses that Sonoran Desert toads synthesize 5-MeO-DMT endogenously or via a microbial symbiont.

## Introduction

Secondary metabolites are bioactive substances that do not directly contribute to an organism’s growth, and are highly variable in both composition and mechanism of action (Torres and Schmidt, 2019; Torres et al., 2020). These small molecules are the targets of intensive biomedical research to treat conditions from chronic pain to cancer (Singla et al., 2021; Fakhri et al., 2022; Rodríguez-Landa et al., 2023). Many secondary metabolites play important roles in ecological processes such as predation and defense (Bücherl et al,, 1968; Brodie, 2009; Ballinger and Perlman, 2017; Nekaris et al., 2020). Animals are thought to acquire chemical defenses through three main routes (Mebs, 2001): (1) taking elemental building blocks from the environment and producing toxins endogenously (e.g. many venomous snakes produce Phospholipase A2 [Kini, 2003]), (2) via symbionts (e.g. rough-skinned newts [*Taricha torosa*] secrete tetrodotoxin produced by a bacteria living on their skin [Vaelli et al., 2020]), and (3) sequestering chemicals from dietary organisms (e.g. dendrobatid poison frogs sequester alkaloids from invertebrate prey [Saporito et al., 2004; O’Connell et al., 2021]). While the source of some molecules used for chemical defense have been established, the origins of psychedelic compounds are comparatively understudied.

The Sonoran Desert toad (*Incilius alvarius*) secretes a mix of indolealkylamine secondary metabolites, including the potent hallucinogenic compound 5-Methoxy-N,N-dimethyltryptamine (5-MeO-DMT) (Musgrave and Cochran, 1929; Hanson and Vial, 1956; Erspamer et al., 1967). These are among the largest toads native to the United States (Holycross et al., 2022) and are known to eat a variety of invertebrates and occasionally vertebrates (Cole, 1962; Stebbins, 2003). *I. alvarius* is found in a variety of biotic communities, primarily in the Sonoran Desert region of Southern Arizona, USA and Sonora, Mexico. Like other bufonids, *I. alvarius* secretes a mix of toxins from its specialized parotoid and other glands when agitated. However, unlike secretions of related species, *I. alvarius* secretions contain high concentrations of 5-MeO-DMT. In fact, *I. alvarius* is the only animal currently known to secrete 5-MeO-DMT, though it is also found in a number of plants (Ermakova et al., 2021). 5-MeO-DMT causes marked behavioral and psychological effects in humans, which led to its popularization as a recreational drug in the 1980s. In recent decades, there has been significant growth in use and demand for *I. alvarius* secretions, resulting in widespread illegal toad poaching (Villa, 2020).

The source of 5-MeO-DMT in Sonoran Desert toads is unknown. It has been suggested that *I. alvarius* might produce 5-MeO-DMT endogenously from 5-Hydroxy-dimethyltryptamine (bufotenine) with a 5-Hydroxyindole-O-methyltransferase (Erspamer et al., 1967; Yu, 2008), but the existence of this enzyme in toad parotoid glands has not been confirmed. Meanwhile, multiple lines of evidence suggest that the diet of *I. alvarius* might be linked to their secretion of 5-MeO-DMT and other compounds used in defense. Savitzky et al. (2012) examined patterns of defensive sequestration across tetrapods and predict that an untested species will sequester defensive compounds if it is (1) from a known sequestering taxon, (2) consumes toxic prey items, and (3) uses passive defense. *I. alvarius* fits all these criteria as a member of the family Bufonidae (Daly et al., 2007; Hantak et al., 2013) which consumes potentially toxic ants and beetles (Bogert & Oliver, 1945; Cole, 1962) and displays a sitting ‘puff’ behavior when approached by predators (Hanson & Vial, 1956).

In this study, we investigate whether *I. alvarius* specializes on chemically defended prey and if such a specialization correlates with presence of 5-MeO-DMT in *I. alvarius* secretions. Relatively few studies (Erspamer et al., 1967; Schwelm et al., 2021) profile the chemicals secreted by *I. alvarius*, and dietary studies conducted over 60 years ago involved small sample sizes (n≤6) (King, 1932; Bogert & Oliver, 1945; Gates, 1957; Cole, 1962). We analyzed the stomach and fecal contents and tested for 5-MeO-DMT and other secondary metabolites in the secretions of *I. alvarius* from several localities comprising multiple habitats in the vicinity of Tucson, Arizona. We also sampled diet from three sympatric anuran species (including two bufonids) and examined compounds in the secretions from both sympatric bufonids for comparison. We predicted that if *I. alvarius* sequester 5-MeO-DMT from a dietary source, we would see a relationship between diet composition and 5-MeO-DMT load, and/or specialization in arthropod taxa that could produce 5-MeO-DMT relative to sympatric anurans not known produce 5-MeO-DMT. To our knowledge, this is the first study to investigate the relationship of diet and 5-MeO-DMT in *I. alvarius* and other sympatric anurans.

## Materials and Methods

### Field Sampling

From August-September 2020, we sampled across five sites near Tucson, Arizona the following species: Sonoran Desert toads (Bufonidae: *Incilius alvarius*, N=29; average mass 191.2g, range 0.5g-400.0g), red-spotted toads (Bufonidae: *Anaxyrus punctatus*, N=7, average mass 16.3g, range 5.6g–28.0g), Great Plains toads (Bufonidae: *Anaxyrus cognatus*, N=4, average mass 71.8g, range 41.0–88.0g) and Couch’s spadefoots (Scaphiopodidae: *Scaphiopus couchii*, N=10, average mass 27.7g, range 20.5–38.5g). As habitat influences prey availability, we characterized the biotic community (Brown, 1994) at each site and identified whether the habitat was predominantly urbanized (N=2) or native (N=3) (Table 1). We also classified habitat as either riparian (at a river or arroyo), semi-riparian (at an ephemeral pool), or non-riparian (away from any surface water).

**Table 1.** 2020 Pima County Anuran Sampling Localities. Ia = *Incilius alvarius*, Ac = *Anaxyrus cognatus*, Ap = *Anaxyrus punctatus*, Sc = *Sceloporus couchii.* Biotic community classifications are based on Brown (1994).

| Location | Habitat<br>(Biotic Community) | Coordinates<br>(Northing,<br>Easting) | Elevation<br>(m) | Ia | Ac | Ap | Sc |
| --- | --- | --- | --- | --- | --- | --- | --- |
| Santa Cruz River east of "A" Mountain | Urbanized Riparian (Sonoran Desertscrub: Arizona Upland Subdivision) | 32.205787,<br>-110.989112 | 716 | 11 | 1 | 2 | 3 |
| "Mesquite Circle Pond" near Santa Cruz River | Urbanized Semi-Riparian (Sonoran Desertscrub: Arizona Upland Subdivision) | 32.191260,<br>-110.985556 | 726 | 6 | 3 | 3 | 3 |
| East side of Green Valley | Native Non-Riparian (Chihuahuan Desertscrub) | 31.740726,<br>-111.010314 | 958 | 1 | 0 | 0 | 0 |
| Pantano wash near Esmond Station Rd | Native Riparian (Sonoran Desertscrub: Arizona Upland Subdivision) | 32.105829,<br>-110.731685 | 901 | 1 | 0 | 0 | 4 |
| Tanque Verde Falls, lower Rincon/Catalina Mountains | Native Riparian (Sonoran Desertscrub: Arizona Upland Subdivision) | 32.257307,<br>-110.650095 | 1011 | 10 | 0 | 2 | 0 |

To find our study species, we walked with flashlights through suitable habitats on rainy or humid nights. We sampled individuals at various life stages, from juveniles to large adults. We collected anurans using nitrile gloves and stored them individually in clean plastic buckets with air and local substrate. We photographed, weighed, and measured lengths of each anuran before collecting secretions and recent prey items. While we recorded sex based on visual markings for *S. couchii*, which exhibit strong sexual dimorphism, we did not sex *I. alvarius* or *Anaxyrus* spp. because the required procedures are more invasive. We did not collect secretions from Couch’s spadefoots because they lack prominent parotoid glands. We were unable to weigh or measure three *I. alvarius*, five *S. couchii* and two *A. punctatus* due to field constraints.

To collect recent prey items we used a nonlethal procedure to flush the anurans’ stomachs, and collected fecal samples from buckets in which the toads were held. We followed methods described by Solé et al. (2005) to prepare a tube-syringe setup with distilled water. We used a mini spatula to open each individual’s mouth, inserted a soft tube through the esophagus into the stomach, emptied the syringe, and then collected flushed stomach contents with a strainer. We stored flushed stomach contents and fecal samples separately in 1 ml of 100% ethanol. We flushed the stomachs of all 50 individuals sampled, and collected fecal samples for 22/50 individuals comprising all four study species (for more details see **Table S1**). Of the 50 samples anurans, eight had empty stomachs and did not produce fecal samples, and a further six provided samples which were not suitable for visual or molecular identification.

We recorded the exact location of each anuran and, after processing, released them at their capture sites. We decontaminated all gear between field sites using a 5-10% bleach solution (Chellman and McKenny, 2018). All fieldwork was completed under Arizona Game and Fish Department Scientific Collecting Licenses (#SP404533 and #SP407194). All animal work was approved by the Institutional Animal Care and Use Committee of Stanford University (Protocol #33849).

### Mass Spectrometry

We analyzed secretions from parotoid and/or leg glands from 26 *I. alvarius*, six *A. punctatus* and three *A. cognatus.* Secretions were collected in the field using kimwipes (Weil and Davis, 1994). We stored secretion-soaked kimwipes in glass vials containing 2-5 ml of 100% methanol at −20°C until further processing. We swabbed the ventrum and dorsum of selected toads using the same method, and repeat-sampled individuals when they produced more secretions than could be held by one kimwipe or glass vial (see **Table S2** in Supplementary Materials for more detail), resulting in 41 samples used in chemical analysis.

Chemical analysis was conducted using a Direct Analysis in Real Time ionization coupled to a time-of-flight mass spectrometer (DART-TOFMS) (AccuTOF-DART by JEOL, USA, Inc., Peabody MA USA). This process consisted of dipping a glass capillary melting point tube into the methanol solvent within the sample vials and holding the glass tube in the helium DART gas stream, which was set to 350 °C. Mass spectra were extracted from the mass-calibrated centroided files and the resulting high resolution ions (m/z) were compared against a database which included molecular components described in the literature. Presumptive assignments were made of compounds that had a threshold greater than 5% of the base peak and masses within 5 mmu of the nominal molecular weight.

To characterize chemical profiles of Sonoran Desert toads and sympatric species, we searched within each sample for the presence of several tryptamines and derivatives recorded by Erspamer et al. (1965), Cei et al. (1968), Uthaug et al. (2019), and Schwelm et al. (2021). We then compared the presence and absence of these compounds across individuals, species and sites.

### Stomach Sample Visual Identification

We processed stomach and fecal contents by sorting specimens in petri dishes. We photographed dietary items with a Lumenera Infinity 2 camera mounted on an Olympus dissecting microscope (SZ40) and returned the samples to 100% ethanol at 4°C for storage. We uploaded many of our photos to the online citizen science platform BugGuide (VanDyk, 2023) for identification assistance. We visually identified nearly all arthropods to order, and further identified many to genus or species, with input from numerous expert contributors (see Acknowledgements). We additionally classified arthropods as native or introduced species. All photos of prey items are available on DataDryad (submission pending acceptance).

### Stomach Sample Molecular Identification

We isolated DNA from arthropods using the DNEasy Blood & Tissue kit (Qiagen, Hilden, Germany) with adjustments to suit arthropod tissues. We cut the arthropods into ∼0.3cm^3^ (lentil-sized) chunks and crushed them with a plastic pestle in a microcentrifuge tube, then incubated samples at 56°C overnight in Proteinase-K and lysis buffer ‘ATL’ from the DNEasy Blood & Tissue kit. We extracted genomic DNA according to the manufacturer’s instructions and stored it at −xs4°C for downstream amplification. We used PCR to amplify a segment of the cytochrome oxidase 1 (*CO1* or *cox1*) gene from mitochondrial DNA, a standard marker for DNA barcoding (Herbert at al., 2003; Pečnikar and Buzan, 2014). We amplified DNA using general arthropod primers mlCOIintF (5’- GGWACWGGWTGAACWGTWTAYCCYCC) and LOBO-R1 (5’-TAAACYTCWGGRTGWCCRAARAAYCA) from (Leray et al., 2013)) and (Lobo et al., 2013)). For all reactions, we used 1.2 μL of each primer (10 μM) and 21.6 μL of 2X Phusion High-Fidelity PCR Master Mix with GC Buffer (New England Biolabs, Ipswich, MA, USA) in a total reaction volume of 40 μL. We used the following PCR program to amplify *CO1*: 98°C for 60s; 5 cycles of 98°C for 10 s, 48°C for 120 s, 72°C for 1 min; 30 rounds of 98°C for 10 s, 54°C for 120 s, and 72°C for 1 min; and a single incubation of 72°C for 5 min. We ran PCR reactions on a 1% SyberSafe/agarose gel (Life Technologies). We purified successful reactions with a single band of the expected size with the E.Z.N.A. Cycle Pure Kit (Omega Bio-Tek, Norcross, GA, USA). Purified PCR products were Sanger sequenced by GeneWiz Inc. (Cambridge, MA, USA) and we uploaded sequences to GenBank (submission pending).

We used DNA barcode sequences to identify the prey items recovered from stomach contents. Barcode sequences were imported into Geneious Prime (v 2021.1.1) for quality trimming and alignment of forward and reverse sequencing reactions. We used nucleotide BLAST from the NCBI Genbank nr database to identify arthropod sequences to the species level. For genomic data we considered results that yielded greater than 98% sequence similarity as sufficient to assign genus or species, a more conservative threshold than previous studies (Hebert et al., 2003; Moskowitz et al., 2018) to better align with our visual identification results. For specimens with less than 98% Genbank similarity, we assigned an order or family only when the top BLAST result matched that of visual identification.

### Identification Summary

We recorded 280 dietary arthropods for 37 of the 50 anurans sampled, as well as many rocks, sand, and plant material. Following DNA extraction we confirmed 166 arthropod samples contained DNA of suitable quality and quantity for downstream molecular work. Samples from multiple (mixed) animals or where PCR amplification failed to produce a band when visualized on agarose gel were excluded from downstream analysis, resulting in 86 samples suitable for Sanger sequencing. We successfully assigned 34 (39.5%) of sequenced samples with a taxonomic ID using this method. Across molecular and visual methods, we identified 236 (84.3%) samples to family, 55 (19.6%) samples to genus, and 34 (12.1%) samples to species. Of our genus and species-level identifications, 7 (12.7%) were the result of Sanger sequencing only, 33 (60%) were the result of visual examination only and 15 (27.3%) were the result of a consensus identification between visual and molecular methods. In only one case did our molecular and visual methods result in conflicting identifications (*Pogonomyrmex barbatus* vs. *P. rugosus*, see Discussion).

### Data Analysis

We performed data analyses using RStudio version “Ghost Orchid” (2021.09.0+351.pro6) and R version 4.1.2. For all data analyses, prey items from stomach and fecal samples were treated together, and samples with zero identifiable arthropod prey items were excluded. We conducted data reduction using Principal Component Analysis (PCA) with the prcomp function in R. For explanatory variables, we considered the number of prey items in each order, but not the number of rocks or plants. We compared *I. alvarius* diet data between urbanized and native habitats and *I. alvarius* diet data to those of sympatric anuran species (pooled due to low sample sizes of each sympatric species individually). We visualized PCAs using the *ade4* package (Dray & Dufour, 2007) and used the aov() function to test for group differences in principal components. To test whether number of hymenopterans, number of beetles, number of native arthropods, and number of introduced arthropods varied between *I. alvarius* in native and urbanized habitats, we used Wilcoxon rank sum tests with continuity correction (wilcox.test in the R base package). We also used Wilcoxon rank sum tests to test whether the number of beetles, hymenopterans, and other arthropods (pooled) varied between *I. alvarius* and other sympatric anurans. We considered adjusting for anuran weight in our analyses, but we did not detect a significant relationship between weight and diet composition in exploratory generalized linear models.

## Results

### 5-MeO-DMT detected in all Sonoran Desert toad secretions

We first confirmed that Sonoran Desert toad secretions contain 5-MeO-DMT using Direct Analysis in Real Time Mass Spectrometry (DART-TOF MS; **Figure 1**). Indeed, 5-MeO-DMT was detected in parotoid and leg gland secretions from all 26 *I. alvarius* individuals. We also recorded the presence of 5-HO-DMT (bufotenine), 5-MeO-N-methyltryptamine, MeO-tryptamine, 5-MeO-tryptophol and tryptophan in the secretions of *Incilius alvarius*. Bufotenine, 5-MeO-N-methyltryptamine, MeO-tryptamine, 5-MeO-tryptophol and tryptophan were also detected in *Anaxyrus cognatus* or *A. punctatus* (**Figure 1, Table S3**). We detected more diversity in tryptamine-derived compounds in *I. alvarius* ventral and dorsal samples than secretions from the parotoid glands (**Table S3**).

**Figure 1.**
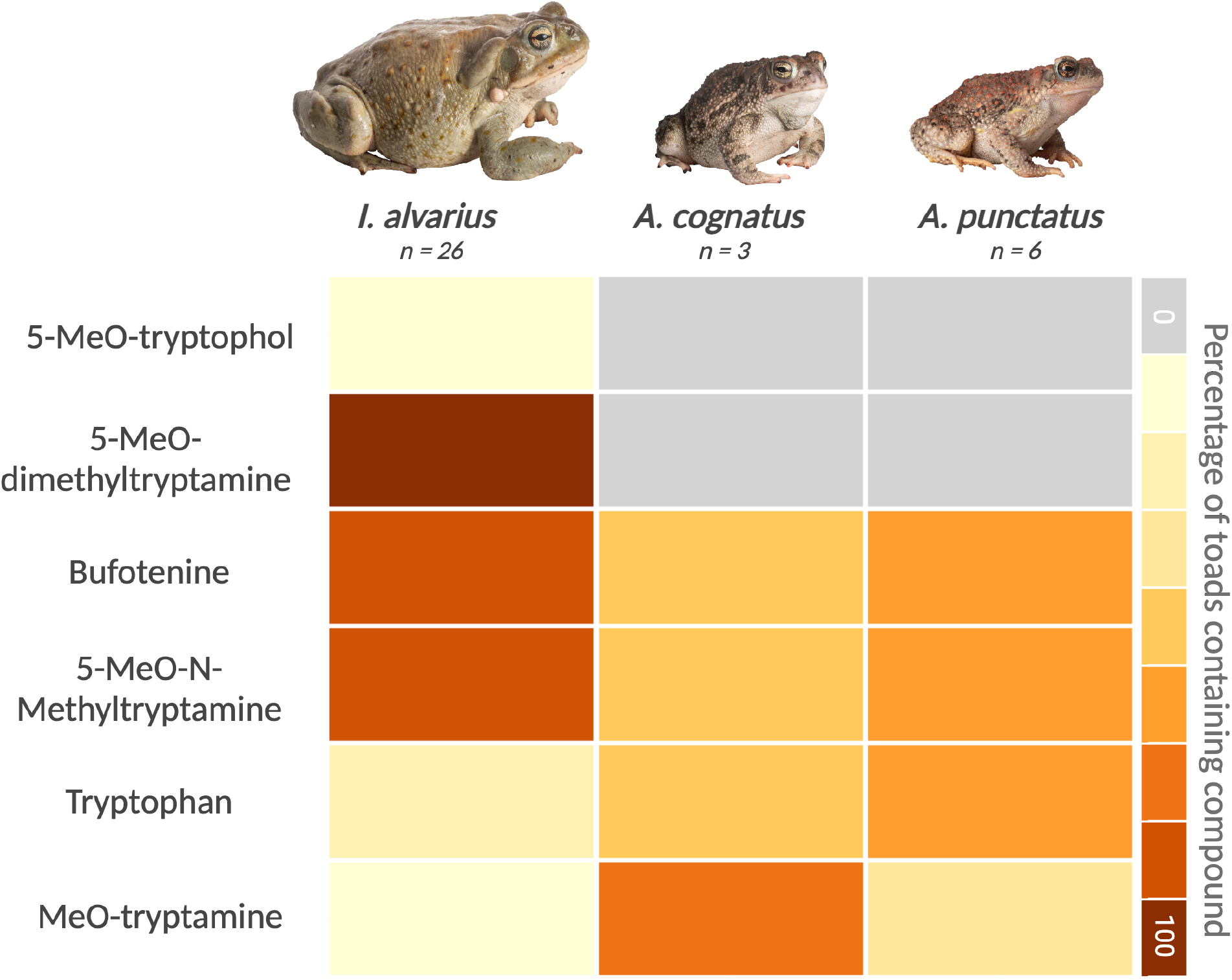
Tryptophan derivatives of interest found in *Anaxyrus cognatus*, *Anaxyrus punctatus* and *Incilius alvarius.* A heatmap indicates the fraction of toads in each species (rows) for which a given compound was detected (columns).

### Sonoran Desert toads are diet generalists

We classified 216 arthropod prey items to seven orders from flushed stomach content and fecal samples of *I. alvarius*. The dominant prey types were hymenopterans and coleopterans (beetles). We found 135 hymenopterans (62.5% of total prey items) in the stomachs of nine *I. alvarius* (33%), and 68 beetles (31.5% of total prey items) in the stomachs of 17 *I*. *alvarius* (56.7%). Of the 135 hymenopterans, 134 were ants. We also found small numbers of Orthoptera, Scorpiones, Blattodea, Araneae and Odonata (**Figure 2; Table S1**).

**Figure 2.**
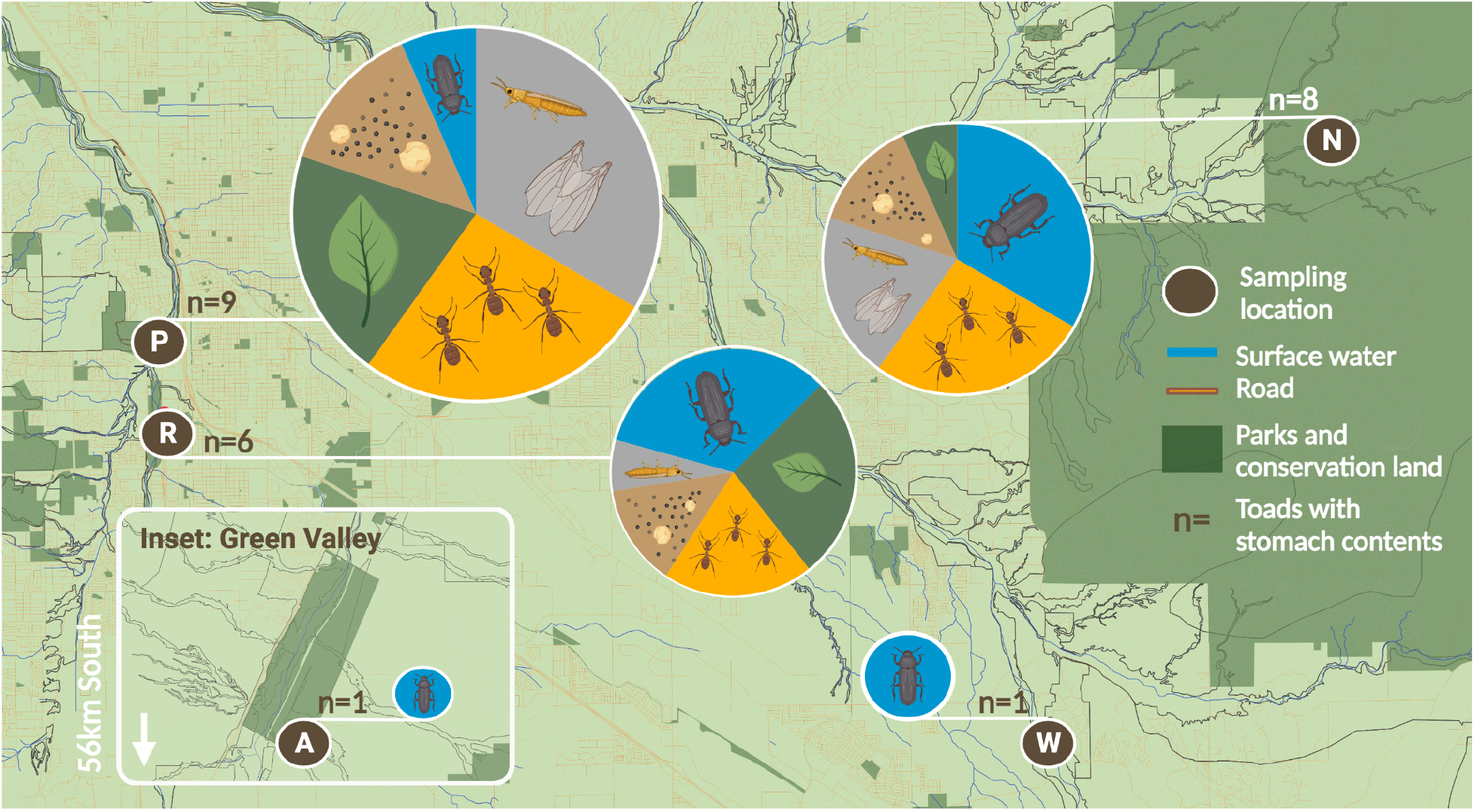
Diet Composition of *I. alvarius* from 5 field sites in Tucson, Arizona. Pie charts represent the average diet composition for all toads at a given site. Diet items are coded in blue = beetles; orange = ants; gray = other arthropods; brown = rocks and sand; green = plant material. Site abbreviations correspond to Table 1: (P) Santa Cruz River east of “A” Mountain; (R) “Mesquite Circle” at Paseo De Las Iglesias near Santa Cruz River; (A) East side of Green Valley; (W) Pantano watershed near Esmond Station Rd; (N) Tanque Verde Falls site in Aliso Canyon, lower Rincon/Catalina Mountains. The baselayer for this diagram was designed in QGIS using data from Pima County (pimamaps.gov) and illustrations created with BioRender.com.

We further classified 185 prey items to 12 families, 36 prey items to 17 genera, and 22 prey items to nine species in *I. alvarius* stomach contents and fecal samples. A full list of identified taxa is available in **Table S4**. Among the prey items, we identified a large wasp (Vespidae, **Figure 3**) and an Arizona bark scorpion (*Centruroides sculpturatus*, **Figure 3**) in the stomach contents of two different adult *I. alvarius*. Both of these species are toxic and capable of delivering formidable stings (Curry et al., 1983; Arif and Williams, 2022).

**Figure 3.**
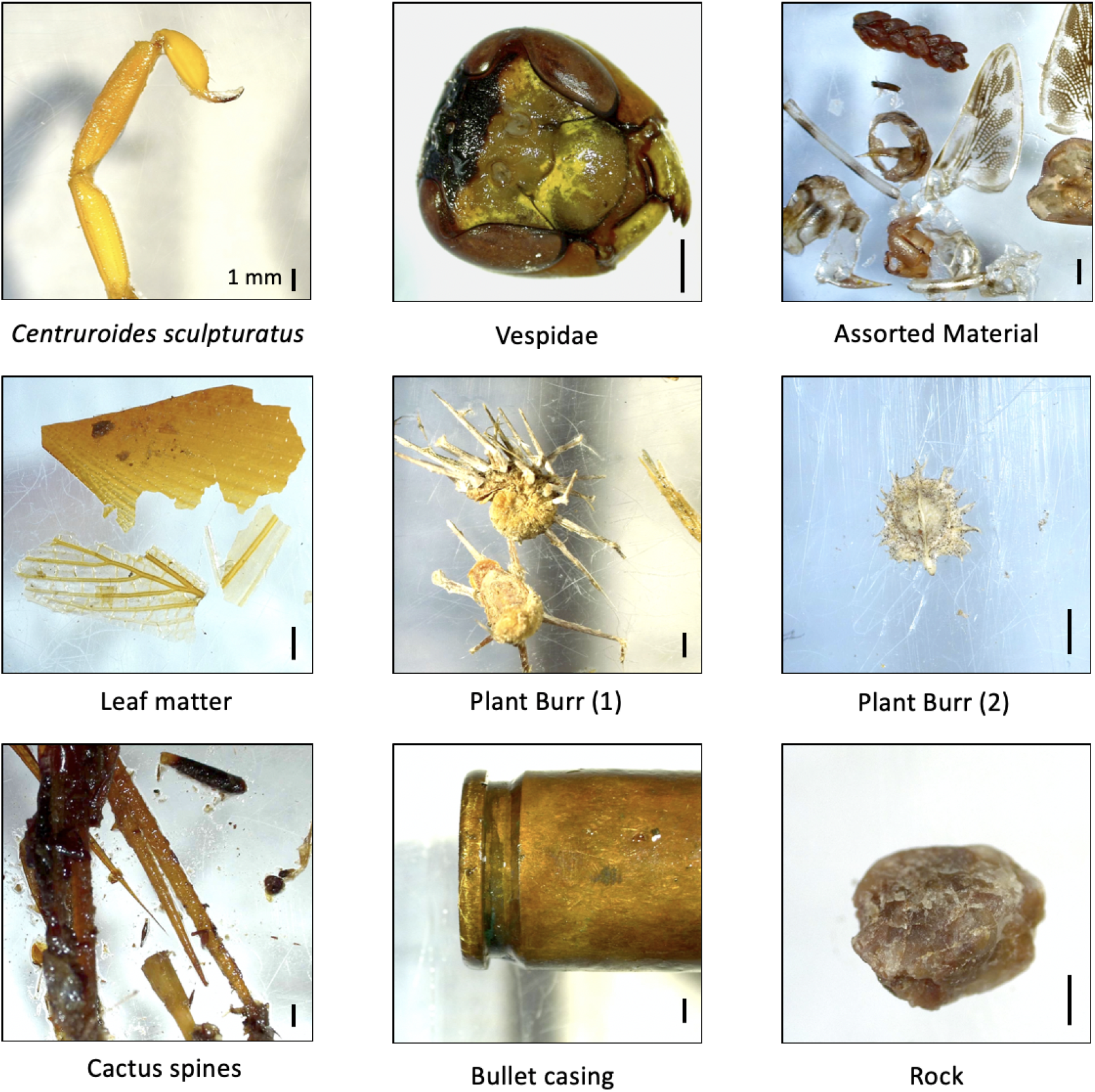
Notable items from *I. alvarius* stomach and fecal contents. All scale bars are 1 mm. Assorted material includes plant matter as well as wings and exoskeleton parts from a dragonfly, perhaps *Pantala flavescens*. The presence of a bullet casing in *I. alvarius* fecal contents is detailed in Luccioni and Wyman (2021).

In addition to arthropod prey, we identified plant matter and inorganic material in the stomach contents of many *I. alvarius*. Of the 30 *I. alvarius* sampled, 13 (43%) had plant matter and 7 (23%) had rocks or sand (gastroliths) in their stomach or fecal contents. Specific items included leaves, stems, burs, cactus spines, and a bullet casing (Luccioni and Wyman, 2021).

### Incilius alvarius in native and urban habitats differ in diet

We first explored the overall patterns of diet in *I. alvarius* using a principal component analysis (PCA) of prey grouped by order (**Figure 4**). Principal component (PC) 1 explained 26.5% of the variance while PC2 explained 20.5%. Toads from different habitats separated significantly by diet in PC2 (F_1_ = 6.21, p = 0.02). We next tested whether specific classes of prey items differed in toads from native and urbanized habitats, but no significant differences were observed in the number of hymenopterans (Wilcoxon, W=54.5, p=0.48), beetles (Wilcoxon, W=92.5, p=0.09), or other arthropods pooled (Wilcoxon, W=40, p=0.08). We identified far more native arthropods than introduced arthropods in Sonoran Desert toad stomach content and fecal samples (211 native, 5 introduced). The five introduced arthropods were only found in toads from urban habitats, and comprised cockroaches of two species (*Periplaneta americana* and *Shelfordella lateralis*).

**Figure 4.**
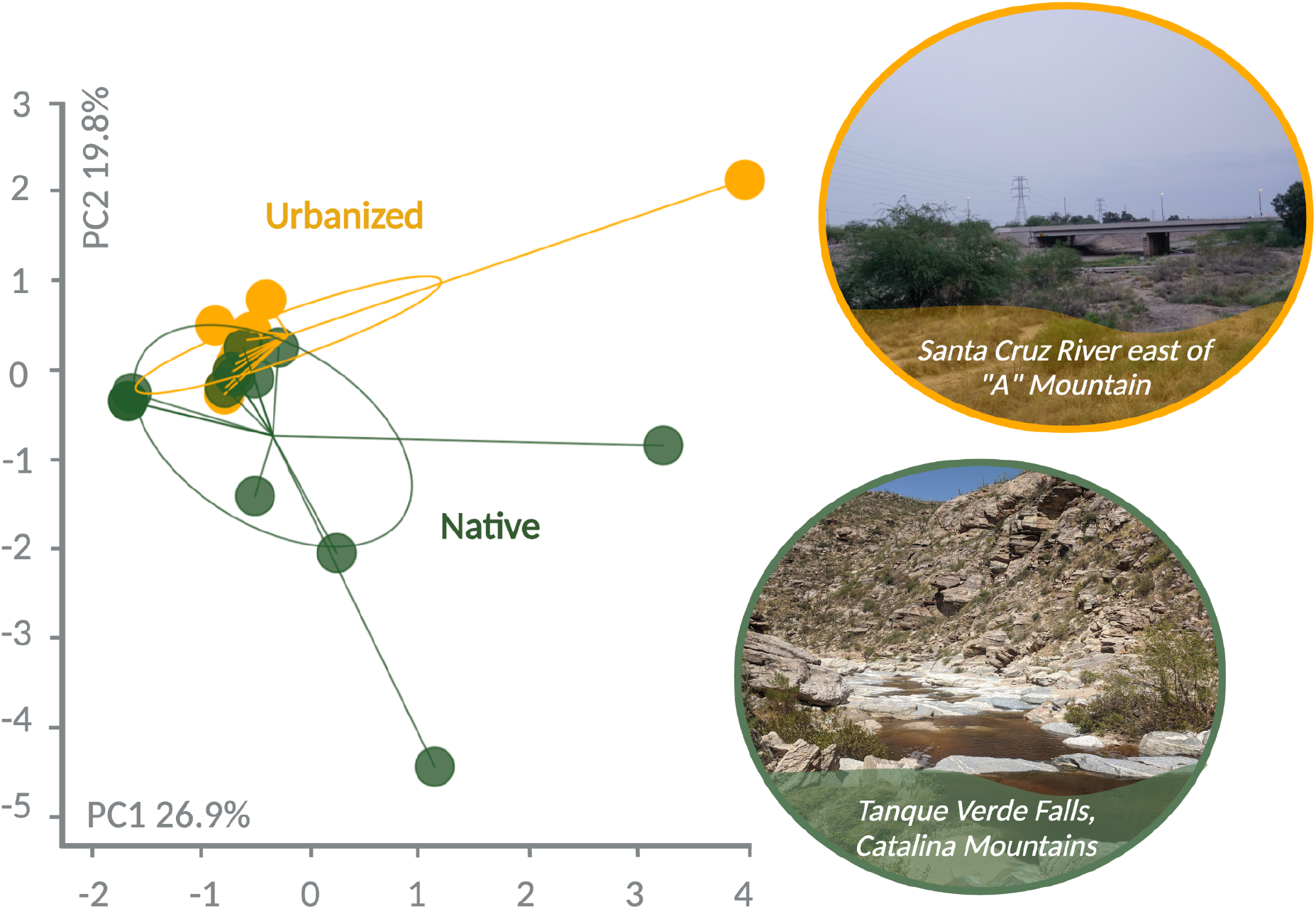
Principal component analysis clustering of Sonoran Desert toad diet data (grouped by order), colored by habitat type (native green, urbanized orange). Each point represents an individual toad; inertia ellipses graphically represent point clouds for each habitat. Photographs depict representative habitat from two of our five study sites classified as urbanized or native.

### Incilius alvarius diet is broadly similar to that of sympatric anurans

To explore whether the *I. alvarius* exhibits dietary specialization, we also sampled the diet of sympatric anurans that do not produce 5-MeO-DMT for comparison. We classified 64 prey items to 8 orders from flushed stomach contents and fecal samples of 7 *A. punctatus*, 4 *A. cognatus*, and 10 *S. couchii* (see **Table S1** for details). We first explored the overall patterns of diet using a PCA of prey grouped by order (**Figure 5**). Principal component (PC) 1 explained 23% of the variance while PC2 explained 18%, although species differences did not explain this variance (p ≥ 0.68 for both principal components). At the level of order, we documented similar diet composition between sympatric anurans and *I. alvarius*, where there were no significant differences in the number of hymenopterans (Wilcoxon, W=136, p=0.41), beetles (Wilcoxon, W=212.5, p=0.10) or other arthropods pooled (Wilcoxon, W=164.5, p=0.91) eaten. Overall, our data support the role of *I. alvarius* and congeners as dietary generalists and opportunists that consume a wide variety of prey.

**Figure 5.**
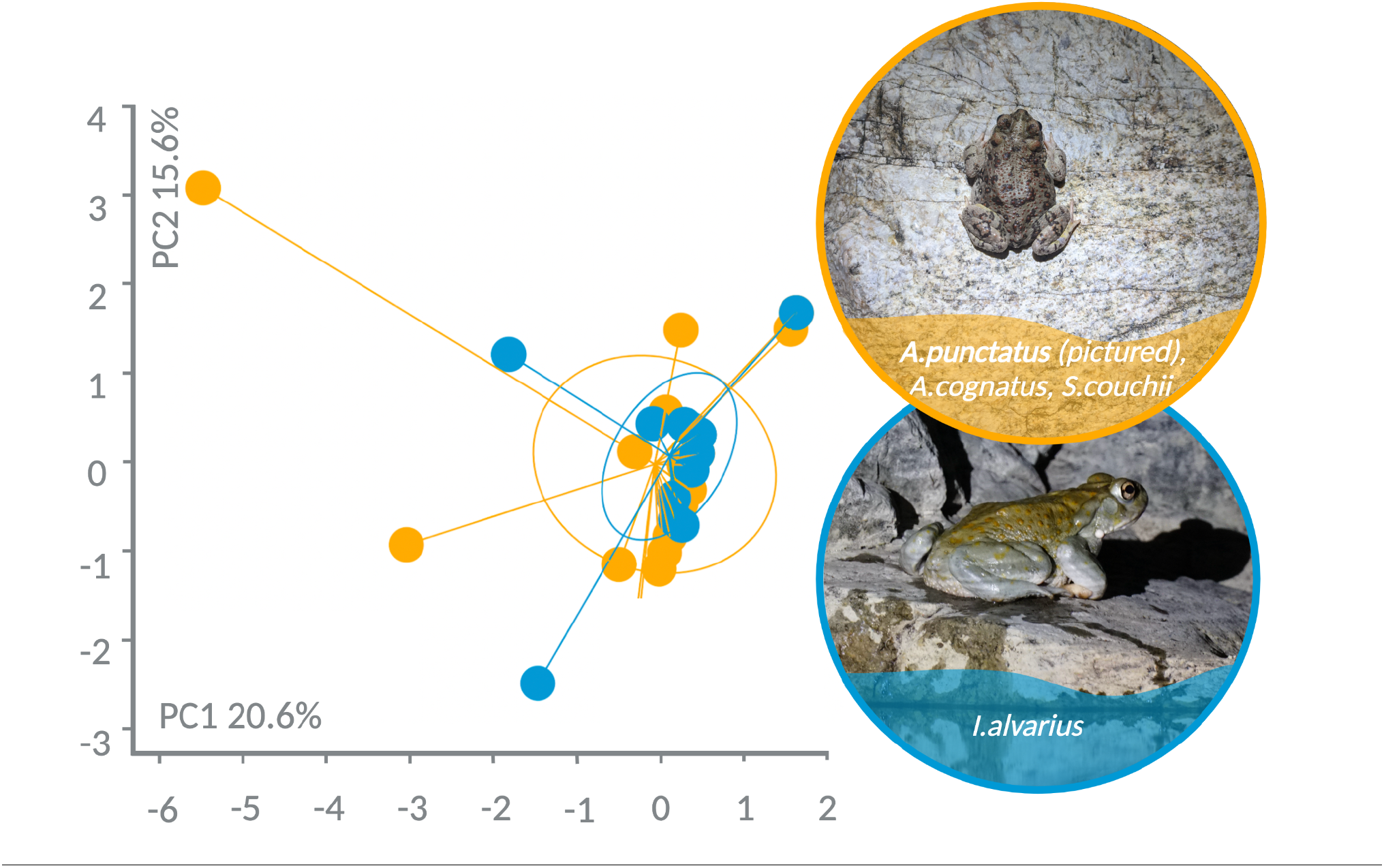
Principal component analysis clustering of anuran diet data (grouped by order), colored by taxon (*I. alvarius* in orange and other species in blue). Each point represents an individual toad; inertia ellipses graphically represent point clouds for each taxon.

## Discussion

### Incilius alvarius diet is similar to sympatric anurans

Animals that sequester chemical defenses tend to demonstrate a preference for toxic prey compared to their non-sequestering relatives (Darst et al., 2005; Yoshida et al., 2020; Moskowitz et al., 2022: bioRxiv 495949). We predicted that if *I. alvarius* obtains 5-MeO-DMT through dietary sequestration, we would observe dietary differences between Sonoran Desert toads and sympatric anurans. Here, we found ants and beetles, many of which possess chemical defenses, to be the dominant prey items for *I. alvarius*. However, we found these groups to be dominant prey items for sympatric anurans as well. Overall, our results are in line with previous dietary studies of this species as well as many other bufonids (Cole, 1962; Núñez et al., 2021). However, we note that sampling stomach contents and fecal samples may overrepresent hard-shelled prey items, which are more likely to remain intact after consumption. Nevertheless, we documented such large numbers of ants and beetles that these two groups likely represent the dominant prey items for Sonoran Desert toads.

Many beetles and ants are chemically defended. *Incilius alvarius* was the only species in our study that we documented consuming *Eleodes* spp. darkling beetles (7 beetles across 3 toads). Although *Eleodes* produce benzoquinones (Blum and Crain, 1961; Happ, 1968; also see Notable Prey below), there is no clear link to production of 5-MeO-DMT or immediate precursors in these taxa, and they may be avoided by (smaller) sympatric anurans simply due to their large size. We also note that several ants of the *Pogonomyrmex* genus were present in our sample set. Indigenous groups including the Kitanemuk, Kawaiisu, Tubatulabal, Interior Chumash, Yawelmani, Wikchamni, Yawdanchi, Bokninwad, Yokod, Palewyami, and possibly the Northern Miwok in California have knowledge and traditions surrounding the ingestion of harvester ants, likely *Pogonomyrmex californicus*, to induce hallucinations (Groark, 1996, 2001). As the symptoms of ingesting *P. californicus* broadly match those of 5-MeO-DMT intoxication from *I. alvarius*, we initially speculated that *I. alvarius* could sequester 5-MeO-DMT from *Pogonomyrmex*. We found three *Pogonomyrmex cf. rugosus* (*P. rugosus* and *P. barbatus* hybridize extensively in the region [Cahan and Keller, 2003], and our molecular identification methods produced variable results) in the stomach contents of two *I. alvarius*. However, we also found several *Pogonomyrmex cf. rugosus* in the stomach contents of two *S. couchii* and one *A*. *cognatus*. Sonoran Desert toads do not appear to specialize on these ants relative to their congeners, nor is there evidence that they eat them in sufficient quantities to provide a major source of 5-MeO-DMT.

Our diet results suggest that a direct dietary source of 5-MeO-DMT is unlikely in Sonoran Desert toads. This is reflected by a lack of prey specialization compared to sympatric species. Additionally, our principal component analysis and documentation of introduced prey in urban habitats only suggest that *I. alvarius* eat somewhat different prey in different habitats, likely reflecting differences in prey availability, but do not demonstrate corresponding differences in 5-MeO-DMT production. To further confirm these results, laboratory studies could vary the diet of captive toads and measure subsequent 5-MeO-DMT production, and/or use mass spectrometry to screen for 5-MeO-DMT in prey items. Repeated sampling of individuals may also be necessary to more fully catalog diet, as our stomach content samples represent a single snapshot in time for each individual.

### Alternatives to dietary sequestration

Several other possibilities exist for Sonoran Desert toad production or acquisition of 5-MeO-DMT. Erspamer et al. (1967) concluded that 5-MeO-DMT is likely formed on the skin of Sonoran Desert toads via the O-methylation of bufotenine (5-HO-DMT), which represents a major component of the secretions from multiple bufonid toads including *I. alvarius*. In light of our results, we consider *in vivo* transformation of 5-MeO-DMT from bufotenine a likely production pathway. Future research should focus on identifying and isolating the hypothesized 5-hydroxyindole-O-methyltransferase necessary for such a transformation.

The possibility of a plant or fungal source for 5-MeO-DMT should not be ruled out because the best-known sources of 5-MeO-DMT are from plants (Ermakova et al., 2022), and multiple other psychedelics are produced by fungi (Guzmán, 2009). Many *I. alvarius* in our study ate plants. In previous literature these occurrences have been attributed to accidental ingestion, but this assumption may represent a “vegetal blind spot” in the diet literature. For example, a relative of *I. alvarius*, *Rhinella arenarum*, has been suspected to behave herbivorously during their dormant period, at least in laboratory conditions (Jungblut et al., 2013). Many of our study anurans had plant matter in their stomach contents and/or items that were too homogenized or fragmented to identify using visual or Sanger sequencing methods, but are likely identifiable with next-generation sequencing. Metabarcoding approaches with a focus on arthropod, plant and fungus-specific primers would help provide a more complete estimation of diet composition for all the anurans in our dataset.

Finally, the skin microbiome may play a role in producing 5-MeO-DMT. Hayes et al. (2009) document the biotransformation of cane toad (*Rhinella marina*) bufadienolides by several bacteria species isolated from parotoid glands and Vaelli et al. (2020) demonstrated the production of tetrodotoxin by bacterial symbionts on the skin of rough-skinned newts (*Taricha granulosa*). Although there is no published research on the microbiome of *I. alvarius*, the roles of these bacteria in related species raise the question of whether skin-dwelling microbial symbionts influence the composition of Sonoran Desert toad secondary metabolites.

### Tryptophan derivatives in the skin and glands of I. alvarius and sympatric bufonids

Toad gland secretions contain a diverse cocktail of secondary metabolites, including various derivatives of tryptophan. In a recent study, Schwelm et al. (2021) explored the toxin composition of five field-collected *I. alvarius* and noted 5-MeO-tryptamine and two positional isomers of hydroxylated MeO-DMT. We additionally recorded, for the first time as far as we are aware, the presence of 5-MeO-tryptamine in *A. cognatus* and 5-MeO-N-methyltryptamine in both *A. cognatus* and *A. punctatus* secretions. We recorded low amounts of 5-MeO-tryptamine and bufotenine for *I. alvarius* and it is likely that the high concentrations of 5-MeO-DMT in remaining samples inhibited detection of other molecules present at lower concentrations. This is supported by our findings that toxin extracts from the ventrum and dorsum, which contain less 5-MeO-DMT, possibly allowed for the detection of more diverse compounds. This inference is supported by the interpretation of the mass spectra of I. *alvarius*, where the high levels 5-MeO-DMT (i.e., the base peak) concealed the compounds below the threshold level of 5%. As Direct Analysis in Real Time (DART) ionization has limited quantitative capabilities, and because we ran variable amounts of toxin from each toad, we were not able to precisely quantify the percent abundance of 5-MeO-DMT in our samples. Thus, we were not able to perform a robust quantitative analysis of variation in 5-MeO-DMT production between toads. In future work, we will use more quantitative approaches to measure 5-MeO-DMT in Sonoran Desert toad excretions.

Bufotenine is a tryptamine alkaloid and an isomer of the psychedelic compound psilocin. We recorded the presence of bufotenine in Sonoran Desert Toads and their congeners. Bufotenine was present in *A. punctatus* at levels >40% but only found in trace amounts (<5%) in *I. alvarius*, which is what we might expect to see if bufotenine is being used as a substrate for creating 5-MeO-DMT. Many additional tryptamines and derivatives reported in Sonoran Desert toad secretions (see Erspamer et al., 1965; Cei et al., 1968, Uthaug et al., 2019; Schwelm et al., 2021) we also found in secretions of *A. cognatus and A. punctatus*. Some of these molecules are potential chemical precursors for 5-MeO-DMT. Understanding the similarities in toxin composition between *Incilius alvarius* and related anurans that likely do not produce 5-MeO-DMT can help pinpoint the ways that *I. alvarius* are unique and provide clues as to which pathways are used to create 5-MeO-DMT.

### I. alvarius as dietary generalists capable of ingesting toxic and barbed prey

Our findings support previous results (Cole, 1962; Gates, 1957) that indicate Sonoran Desert toads are largely dietary generalists and opportunists, observed to consume a wide variety of prey, including arthropods with stings or toxic secretions (Bogan & Eppehimer, 2017). However, they may exhibit some specialization on ants and beetles, as these two groups made up the vast majority of dietary prey items, which was also documented by Cole (1962). In addition to many ants and beetles, we also found spiders, cockroaches, dragonfly larvae, a grasshopper, a cricket, a wasp, and a scorpion in Sonoran Desert toad stomach contents. Unlike previous studies, we did not document vertebrate prey in the stomach contents of any Sonoran Desert toad sampled.

In line with Cole (1962), we recorded a variety of arthropods with stings or toxic secretions consumed by *I. alvarius.* These included putative rough harvester ants (*Pogonomyrmex cf. rugosus*) and a vespid wasp, both of which readily sting and possess potent venoms (Pinnas et al., 1977; Arif and Williams, 2022), as well as a variety of *Eleodes* and other tenebrionid beetles, many of which produce a noxious secretion containing benzoquinones when disturbed (Blum & Crain, 1961; Happ, 1968). Perhaps most remarkable was the tail of an Arizona bark scorpion (*Centruroides sculpturatus*), whose sting is extremely painful and potentially lethal to humans (Curry et al., 1983), and produces neurotoxic effects in the nerve fibers of at least one species of frog (Cahalan, 1975). It is unknown how Sonoran Desert toads and their relatives (we also found *Pogonomyrmex* in *A. cognatus* and *S. couchii* stomach contents) manage to consume arthropods with such formidable chemical defenses. Horned lizards (*Phrynosoma* spp.) have been shown to detoxify *Pogonomyrmex* ant venom via a blood factor (Schmidt et al., 1989), while southern grasshopper mice (*Onychomys torridus*) are resistant to bark scorpion venom in part due to a mutation in a non-target sodium channel (Rowe et al., 2013). It is possible that *I. alvarius* and other anurans employ similar mechanisms.

In addition to arthropod prey, we identified plant matter and inorganic material in the stomach contents of many *I. alvarius*. These findings are usually explained as the result of incidental capture, either as a byproduct when attempting to capture something else or the result of misidentification. However, gastroliths (stomach stones) are believed to aid in the mechanical breakdown of organic matter in the gut of multiple other taxa (Wings, 2007), raising the possibility that Sonoran Desert toads consume hard inorganic material for digestive purposes.

## Conclusion

We found limited evidence to support the hypothesis that *Incilius alvarius* 5-MeO-DMT production is linked to diet. Instead, we found that *I. alvarius* from multiple sites and habitats all produce high quantities of 5-MeO-DMT, despite a diversity of prey in their stomach and fecal contents. Furthermore, we found that *I. alvarius* have a largely similar diet to three sympatric anuran species which likely do not produce 5-MeO-DMT. Although our study was limited to single diet items, our results do not support a direct dietary arthropod source of 5-MeO-DMT in *I. alvarius*. Alternatively, these toads could produce 5-MeO-DMT endogenously using a 5-hydroxyindole-O-methyltransferase or via a bacterial symbiont, as previously hypothesized, or sequester 5-MeO-DMT from a plant or fungal source.

## Acknowledgements

We are grateful to a wonderful team who made this study possible: Leisy and Michael Wyman allowed us facilities access and indispensable fieldwork support; Robert Villa shared his expertise and advice regarding wide-ranging aspects of the study; Professor John Weins at the University of Arizona stored our samples from August to September 2020; Dr. Nora Martin advised our stomach flushing protocol; Dave Ramirez provided valuable logistical support and Katie Fiocca shared helpful comments on our manuscript. Many people generously assisted with visual identification of invertebrates from stomach and fecal samples, including Brad Barnd (BugGuide), Dr. Michael Bogan, Dr. David Donoso, Curt Harden (BugGuide), Solomon Hendrix (BugGuide), Andrew Johnston, Blaine Mathison (BugGuide) and Wyatt Mendez. We also thank Alberto Arevalo and Elizabeth Navarro for their support in creating photographs for select figures, and Quinn Agnew, Daniel Carhuff, Andrew Connoy and Tasman Rosenfeld for their assistance with our fieldwork.

## Appendices

Table S1. Summary of Dietary Items and Flushed and Fecal Samples from *Incilius alvarius* and Sympatric Species

**Table S2.** Description of Samples for Chemical Analysis.

| Toad Species | Sample Treatment | Number of Toads |
| --- | --- | --- |
| *I. alvarius* | Sampled once according to procedure outlined in Methods | 39 |
| *I. alvarius* | Sampled twice due to large size of parotoid gland | 6 |
| *I. alvarius* | Sampled twice due to suspected contamination from blood spotting as an accidental result of toxin extraction. | 4 |
| *I. alvarius* | Sampled from the parotoid gland, leg gland, dorsal skin and ventral skin. | 1 |
| *I. alvarius* | Sampled from the parotoid gland, dorsal skin and ventral skin. | 1 |
| *A. punctatus* | Sampled from the parotoid gland | 6 |
| *A. cognatus* | Sampled from the parotoid gland | 3 |

**Table S3.**
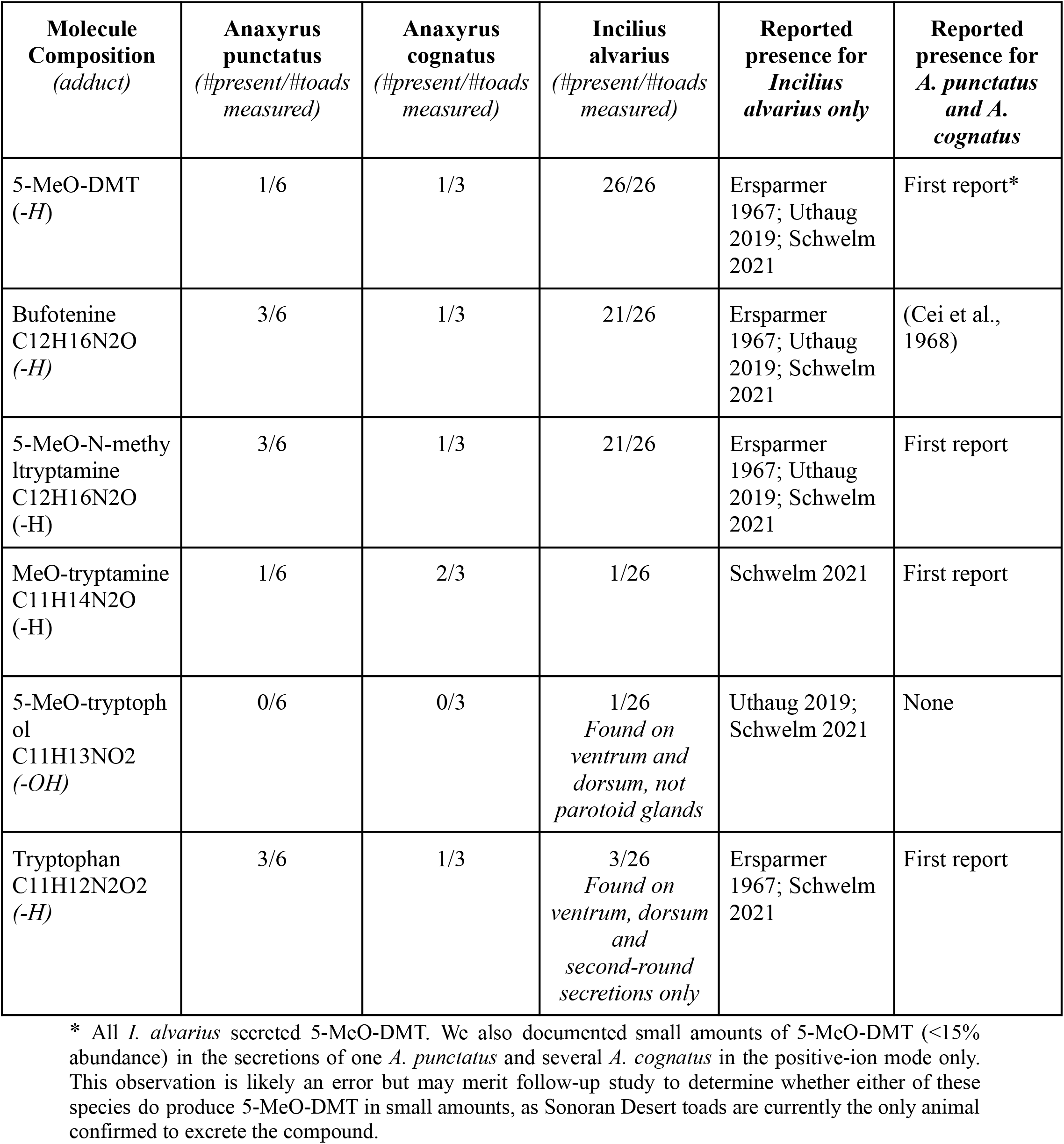
Tryptophan and Derivatives from the Parotoid Glands and Skin of Sampled Anurans.

Table S4. Identifications of arthropod prey consumed by *I. alvarius* and sympatric species. Includes all prey items identified to family or beyond, except ants, which comprised all hymenopteran specimens but one.

**Figure S1.**
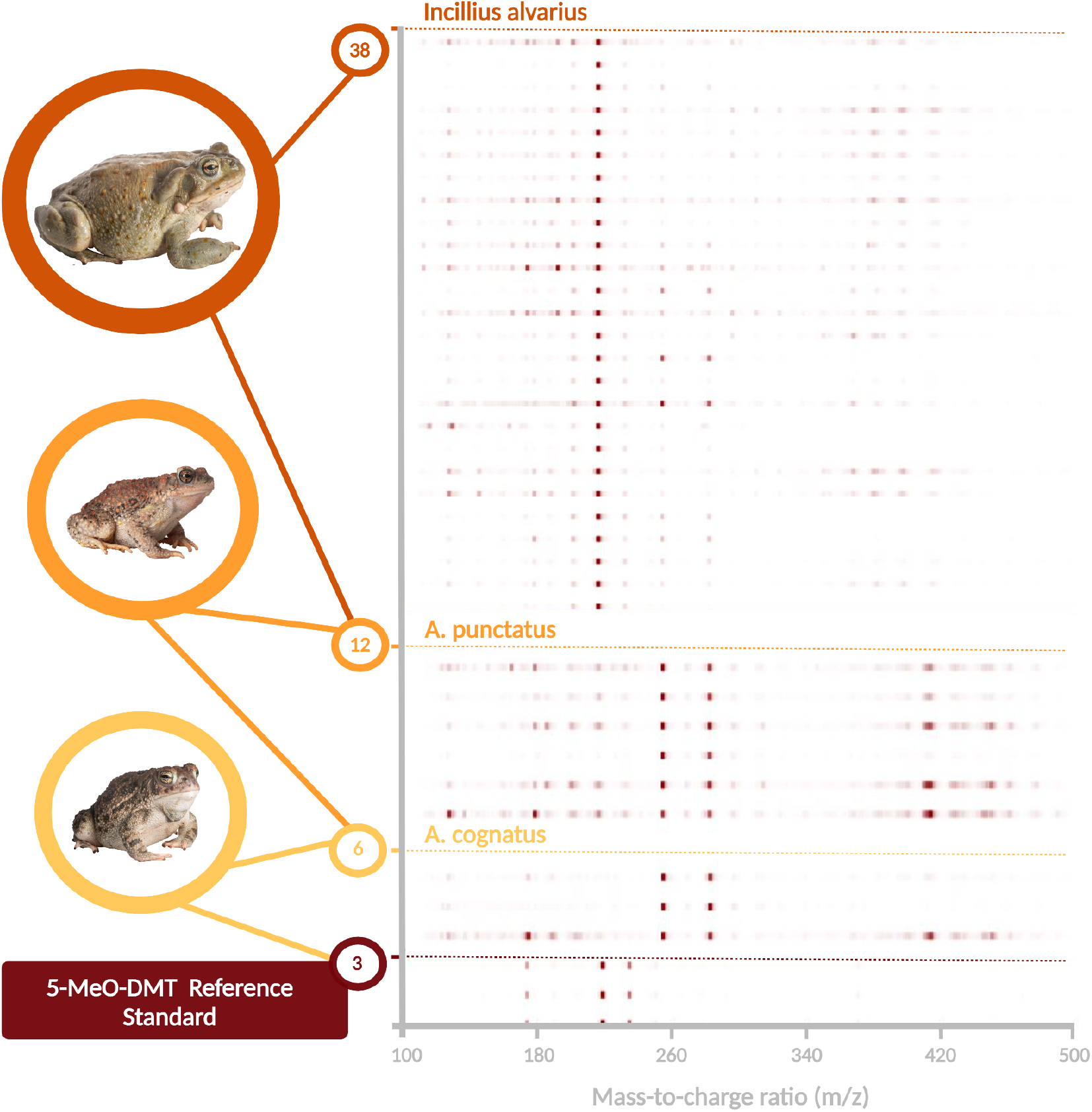
Presence and absence of various molecules across species sampled, as determined by Direct Analysis in Real Time Mass Spectrometry (DART-TOF MS) in the negative ion mode.

### Additional Description of Diet Data for Sympatric Anurans

As in Sonoran Desert toads, hymenopterans and coleopterans (beetles) were the dominant prey items, with 37 ants (57.8%) in the stomach contents of 9 toads, and 19 beetles (29.7%) in the stomach contents of 7 toads (for species-specific prey type breakdowns, see Table M in Supplementary Materials). We encountered 2 introduced black webspinners (*Oligotoma nigra*) in the stomach of a Couch’s spadefoot from urban habitat; all other prey items identified were native species. We found plant matter in the stomach contents of at least one individual of all three species, and rocks or sand in the stomach contents of one Great Plains toad and one Couch’s spadefoot.

## Literature Cited

1. Arif, F., and M. Williams. 2022. Hymenoptera Stings. StatPearls. https://www.ncbi.nlm.nih.gov/books/NBK518972/

2. Ballinger, M. J., and S.J. Perlman. 2017. Generality of Toxins in Defensive Symbiosis: Ribosome-Inactivating Proteins and Defense against Parasitic Wasps in Drosophila. Edited by Greg Hurst. PLOS Pathogens 13 (7): e1006431. 10.1371/journal.ppat.1006431.

3. Becker, Matthew H., and Brian Gratwicke. 2017. “Minimum Lethal Concentration of Sodium Hypochlorite for the Amphibian Pathogen Batrachochytrium Dendrobatidis.” Edited by Tom Coenye. PLOS ONE 12 (4): e0176439. 10.1371/journal.pone.0176439.

4. Blum, Murray S., and Ronald D. Crain. 1961. “The Occurrence of Para-Quinones in the Abdominal Secretion of Eleodes Hispilabris (Coleoptera: Tenebrionidae).” Annals of the Entomological Society of America 54 (4): 474–77. 10.1093/aesa/54.4.474.

5. Bogan, Michael T., and Drew E. Eppehimer. 2017. “Attempted Predation of Western Desert Tarantula By Sonoran Desert Toad.” The Southwestern Naturalist 62 (2): 146–48. 10.1894/0038-4909-62.2.146.

6. Bogert, C. M., and J.A. Oliver. 1945. A Preliminary Analysis of the Herpetofauna of Sonora. American Museum of Natural History, New York, New York.

7. Brodie, Edmund D. 2009. “Toxins and Venoms.” Current Biology 19 (20): R931–35. 10.1016/j.cub.2009.08.011.

8. Brown, D.E. 1994. Biotic communities: southwestern United States and northwestern Mexico. University of Utah Press, Salt Lake City, Utah.

9. Bücherl, W.; E.E. Buckley; and V. Deulofeu. (Eds.). 1968. Venomous animals and their venoms. Academic Press, New York, New York.

10. Cahalan, M D. 1975. “Modification of Sodium Channel Gating in Frog Myelinated Nerve Fibres by Centruroides Sculpturatus Scorpion Venom.” The Journal of Physiology 244 (2): 511–34. 10.1113/jphysiol.1975.sp010810.

11. Cei, J. M., V. Erspamer, and M. Roseghini. 1968. “Taxonomic and Evolutionary Significance of Biogenic Amines and Polypeptides in Amphibian Skin. II. Toads of the Genera Bufo and Melanophryniscus.” Systematic Zoology 17 (3): 232–45. 10.2307/2412002.

12. Chellman, I., and H. McKenny. 2018. Interagency Conservation Strategy for Mountain Yellow-legged Frogs in the Sierra Nevada, Attachment 4: Equipment Decontamination Protocol. National Park Service.

13. Cole, Charles J. 1962. “Notes on the Distribution and Food Habits of Bufo Alvarius at the Eastern Edge of Its Range.” Herpetologica 18 (3): 172–75.

14. Curry, Steven C., Michael V. Vance, Patricia J. Ryan, Donald B. Kunkel, and William T. Northey. 1983. “Envenomation by the Scorpion Centruroides Sculpturatus.” Journal of Toxicology: Clinical Toxicology 21 (4–5): 417–49. 10.3109/15563658308990433.

15. Daly, J. W., J. M. Wilham, T. F. Spande, H. M. Garraffo, R. R. Gil, G. L. Silva, and M. Vaira. 2007. “Alkaloids in Bufonid Toads (Melanophryniscus): Temporal and Geographic Determinants for Two Argentinian Species.” Journal of Chemical Ecology 33 (4): 871–87. 10.1007/s10886-007-9261-x.

16. Darst, Catherine R., Pablo A. Menéndez-Guerrero, Luis A. Coloma, and David C. Cannatella. 2005. “Evolution of Dietary Specialization and Chemical Defense in Poison Frogs (Dendrobatidae): A Comparative Analysis.” The American Naturalist 165 (1): 56–69. 10.1086/426599.

17. Dray, Stéphane, and Anne-Béatrice Dufour. 2007. “The Ade4 Package: Implementing the Duality Diagram for Ecologists.” Journal of Statistical Software 22 (4). 10.18637/jss.v022.i04.

18. Ermakova, Anna O, Fiona Dunbar, James Rucker, and Matthew W Johnson. 2022. “A Narrative Synthesis of Research with 5-MeO-DMT.” Journal of Psychopharmacology 36 (3): 273–94. 10.1177/02698811211050543.

19. Erspamer, V., T. Vitali, M. Roseghini, and J. M. Cei. 1965. “5-Methoxy- and 5-Hydroxy-Indolealkylamines in the Skin ofBufo Alvarius.” Experientia 21 (9): 504–504. 10.1007/BF02138956.

20. Erspamer, V., T. Vitali, M. Roseghini, and J.M. Cei. 1967. “5-Methoxy- and 5-Hydroxyindoles in the Skin of Bufo Alvarius.” Biochemical Pharmacology 16 (7): 1149–64. 10.1016/0006-2952(67)90147-5.

21. Fakhri, S., S.Z. Moradi, M.H. Farzaei, and A. Bishayee. 2022. Modulation of dysregulated cancer metabolism by plant secondary metabolites: A mechanistic review, p. 276-305. In: Seminars in cancer biology. Academic Press.

22. Fišer Pečnikar, Živa, and Elena V. Buzan. 2014. “20 Years since the Introduction of DNA Barcoding: From Theory to Application.” Journal of Applied Genetics 55 (1): 43–52. 10.1007/s13353-013-0180-y.

23. Gates, Gerald O. 1957. “A Study of the Herpetofauna in the Vicinity of Wickenburg, Maricopa County, Arizona.” Transactions of the Kansas Academy of Science (1903-) 60 (4): 403–18. 10.2307/3626396.

24. Groark, K.P. 1996. “Ritual and Therapeutic Use of ‘Hallucinogenic’ Harvester Ants (Pogonomyrmex) in Native South-Central California.” Journal of Ethnobiology 16 (1): 1–29.

25. Groark, K.P. 2001. Taxonomic Identity of ‘Hallucinogenic’ Harvester Ant (Pogonomyrmex californicus) Confirmed. Journal of Ethnobiology 21 (2): 133–144.

26. Guzmán, G. 2009. The hallucinogenic mushrooms: diversity, traditions, use and abuse with special reference to the genus Psilocybe. In: Fungi from Different Environments. J.K. Misra, and S.K. Deshmukh (eds.). CRC Press, Boca Raton, Florida.

27. Hanson, Joe A., and James L. Vial. 1956. “Defensive Behavior and Effects of Toxins in Bufo Alvarius.” Herpetologica 12 (2): 141–49.

28. Hantak, Maggie M., Taran Grant, Sherri Reinsch, Dale Mcginnity, Marjorie Loring, Naoki Toyooka, and Ralph A. Saporito. 2013. “Dietary Alkaloid Sequestration in a Poison Frog: An Experimental Test of Alkaloid Uptake in Melanophryniscus Stelzneri (Bufonidae).” Journal of Chemical Ecology 39 (11–12): 1400–1406. 10.1007/s10886-013-0361-5.

29. Happ, George M. 1968. “Quinone and Hydrocarbon Production in the Defensive Glands of Eleodes Longicollis and Tribolium Castaneum (Coleoptera, Tenebrionidae).” Journal of Insect Physiology 14 (12): 1821–37. 10.1016/0022-1910(68)90214-X.

30. Hayes, R. Andrew, Andrew M. Piggott, Kristian Dalle, and Robert J. Capon. 2009. “Microbial Biotransformation as a Source of Chemical Diversity in Cane Toad Steroid Toxins.” Bioorganic & Medicinal Chemistry Letters 19 (6): 1790–92. 10.1016/j.bmcl.2009.01.064.

31. Hebert, Paul D.N., Sujeevan Ratnasingham, and Jeremy R. De Waard. 2003. “Barcoding Animal Life: Cytochrome c Oxidase Subunit 1 Divergences among Closely Related Species.” Proceedings of the Royal Society of London. Series B: Biological Sciences 270 (suppl_1). 10.1098/rsbl.2003.0025.

32. Helms Cahan, Sara, and Laurent Keller. 2003. “Complex Hybrid Origin of Genetic Caste Determination in Harvester Ants.” Nature 424 (6946): 306–9. 10.1038/nature01744.

33. Holycross, Andrew T., Thomas C. Brennan, and Randall D. Babb. 2022. A field guide to amphibians and reptiles in Arizona. Arizona Game and Fish Department.

34. Jungblut, Lucas David, Andrea Gabriela Pozzi, and Dante Agustín Paz. 2013. “A Curious Case of Herbivory in the Common Toad Rhinella Arenarum Arenarum during Hibernation in Captivity Conditions.” Cuadernos de Herpetología 27 (2): 161–62.

35. King, F. Willis. 1932. “Herpetological Records and Notes from the Vicinity of Tucson, Arizona, July and August, 1930.” Copeia 1932 (4): 175–77. 10.2307/1437256.

36. Luccioni, M., and J. Wyman. 2021. Incilius alvarius (Sonoran Desert Toad). Ingestion of Bullet Casing. Herpetological Review 52 (4): 827.

37. Manjunatha Kini, R. 2003. “Excitement Ahead: Structure, Function and Mechanism of Snake Venom Phospholipase A2 Enzymes.” Toxicon 42 (8): 827–40. 10.1016/j.toxicon.2003.11.002.

38. Mebs, Dietrich. 2001. “Toxicity in Animals. Trends in Evolution?” Toxicon 39 (1): 87–96. 10.1016/S0041-0101(00)00155-0.

39. Musgrave, M. E., and Doris M. Cochran. 1929. “Bufo Alvarius, a Poisonous Toad.” Copeia, no. 173: 96–99. 10.2307/1437192.

40. Nekaris, K.A.I., Marco Campera, Vincent Nijman, Hélène Birot, Eva Johanna Rode-Margono, Bryan Grieg Fry, Ariana Weldon, Wirdateti Wirdateti, and Muhammad Ali Imron. 2020. “Slow Lorises Use Venom as a Weapon in Intraspecific Competition.” Current Biology 30 (20): R1252–53. 10.1016/j.cub.2020.08.084.

41. Núñez, K.; Duré, M.; Zárate, G.; Ortiz, F.; and Mendoza, M. 2021. “Diet of Melanophryniscus paraguayensis (Anura: Bufonidae): an Endemic Species to Paraguay.” Herpetological Conservation and Biology 16 (2): 251–258.

42. O’Connell, L.A., LS50: Integrated Science Laboratory Course, Jeremy D. O’Connell, Joao A. Paulo, Sunia A. Trauger, Steven P. Gygi, and Andrew W. Murray. 2021. “Rapid Toxin Sequestration Modifies Poison Frog Physiology.” Journal of Experimental Biology 224 (3): jeb230342. 10.1242/jeb.230342.

43. Pinnas, J, R Strunk, T Wang, and H Thompson. 1977. “Harvester Ant Sensitivity: In Vitro and in Vivo Studies Using Whole Body Extracts and Venom.” Journal of Allergy and Clinical Immunology 59 (1): 10–16. 10.1016/0091-6749(77)90170-1.

44. Rodríguez-Landa, J.F., D. Scuteri, and L. Martínez-Mota. 2023. Plant secondary metabolites: Potential therapeutic implications in neuropsychiatric disorders. Frontiers in Behavioral Neuroscience 17: 1153296.

45. Rowe, Ashlee H., Yucheng Xiao, Matthew P. Rowe, Theodore R. Cummins, and Harold H. Zakon. 2013. “Voltage-Gated Sodium Channel in Grasshopper Mice Defends Against Bark Scorpion Toxin.” Science 342 (6157): 441–46. 10.1126/science.1236451.

46. Saporito, Ralph A., H. Martin Garraffo, Maureen A. Donnelly, Adam L. Edwards, John T. Longino, and John W. Daly. 2004. “Formicine Ants: An Arthropod Source for the Pumiliotoxin Alkaloids of Dendrobatid Poison Frogs.” Proceedings of the National Academy of Sciences 101 (21): 8045–50. 10.1073/pnas.0402365101.

47. Savitzky, Alan H., Akira Mori, Deborah A. Hutchinson, Ralph A. Saporito, Gordon M. Burghardt, Harvey B. Lillywhite, and Jerrold Meinwald. 2012. “Sequestered Defensive Toxins in Tetrapod Vertebrates: Principles, Patterns, and Prospects for Future Studies.” Chemoecology 22 (3): 141–58. 10.1007/s00049-012-0112-z.

48. Schmidt, Patricia J., Wade C. Sherbrooke, and Justin O. Schmidt. 1989. “The Detoxification of Ant (Pogonomyrmex) Venom by a Blood Factor in Horned Lizards (Phrynosoma).” Copeia 1989 (3): 603–7. 10.2307/1445486.

49. Schwelm, Hannes M, Nicole Zimmermann, Tobias Scholl, Johannes Penner, Amy Autret, Volker Auwärter, and Merja A Neukamm. 2022. “Qualitative and Quantitative Analysis of Tryptamines in the Poison of Incilius Alvarius (Amphibia: Bufonidae).” Journal of Analytical Toxicology 46 (5): 540–48. 10.1093/jat/bkab038.

50. Singla, R. K., A.G. Guimarães, and G. Zengin. 2021. Application of plant secondary metabolites to pain neuromodulation. Frontiers in Pharmacology 11: 623399.

51. Stebbins, R.C. 2003. A Field Guide to Western Reptiles and Amphibians. Third edition. Houghton Mifflin Harcourt, Boston, Massachusetts.

52. Solé, Mirco, Olaf Beckmann, Birgit Pelz, Axel Kwet, and Wolf Engels. 2005. “Stomach-Flushing for Diet Analysis in Anurans: An Improved Protocol Evaluated in a Case Study in Araucaria Forests, Southern Brazil*.” Studies on Neotropical Fauna and Environment 40 (1): 23–28. 10.1080/01650520400025704.

53. Torres, Joshua P., Zhenjian Lin, Jaclyn M. Winter, Patrick J. Krug, and Eric W. Schmidt. 2020. “Animal Biosynthesis of Complex Polyketides in a Photosynthetic Partnership.” Nature Communications 11 (1): 2882. 10.1038/s41467-020-16376-5.

54. Torres, Joshua P., and Eric W. Schmidt. 2019. “The Biosynthetic Diversity of the Animal World.” Journal of Biological Chemistry 294 (46): 17684–92. 10.1074/jbc.REV119.006130.

55. Uthaug, M. V., R. Lancelotta, K. Van Oorsouw, K. P. C. Kuypers, N. Mason, J. Rak, A. Šuláková, et al. 2019. “A Single Inhalation of Vapor from Dried Toad Secretion Containing 5-Methoxy-N,N-Dimethyltryptamine (5-MeO-DMT) in a Naturalistic Setting Is Related to Sustained Enhancement of Satisfaction with Life, Mindfulness-Related Capacities, and a Decrement of Psychopathological Symptoms.” Psychopharmacology 236 (9): 2653–66. 10.1007/s00213-019-05236-w.

56. Vaelli, Patric M, Kevin R Theis, Janet E Williams, Lauren A O’Connell, James A Foster, and Heather L Eisthen. 2020. “The Skin Microbiome Facilitates Adaptive Tetrodotoxin Production in Poisonous Newts.” eLife 9 (April): e53898. 10.7554/eLife.53898.

57. VanDyk, John, ed. 2023. BugGuide.Net: Identification, Images, & Information For Insects, Spiders & Their Kin For the United States & Canada. Iowa State University. https://bugguide.net/. (Accessed 4 October 2023)

58. Villa, R. 2020. Toad Smoke: (Un)Natural History of the Sonoran Desert Toad [Youtube Video]. Tucson Herpetological Society. https://www.youtube.com/watch?v=h71r4cHh3p8

59. Weil, Andrew T., and Wade Davis. 1994. “Bufo Alvarius: A Potent Hallucinogen of Animal Origin.” Journal of Ethnopharmacology 41 (1–2): 1–8. 10.1016/0378-8741(94)90051-5.

60. Wings, O. 2007. “A review of gastrolith function with implications for fossil vertebrates and a revised classification”. Acta Palaeontologica Polonica 52 (1).

61. Yoshida, Tatsuya, Rinako Ujiie, Alan H. Savitzky, Teppei Jono, Takato Inoue, Naoko Yoshinaga, Shunsuke Aburaya, et al. 2020. “Dramatic Dietary Shift Maintains Sequestered Toxins in Chemically Defended Snakes.” Proceedings of the National Academy of Sciences 117 (11): 5964–69. 10.1073/pnas.1919065117.

62. Yu, Ai-Ming. 2008. “Indolealkylamines: Biotransformations and Potential Drug–Drug Interactions.” The AAPS Journal 10 (2): 242. 10.1208/s12248-008-9028-5.

