## Supplemental Table 1 for "Diet and chemical defenses of the Sonoran Desert toads"

| Identification Number | Date | Species | Snout-Vent Length (cm) | Weight (g) | Sex (if known) | Age Class | Site | Habitat | Sample Type | Number of Dietary Items | | | | | | | | | | | | | |
|---|---|---|---|---|---|---|---|---|---|---|---|---|---|---|---|---|---|---|---|---|---|---|---|
| | | | | | | | | | | Coleoptera | Odonata | Orthoptera | Embioptera | Hymenoptera | Scorpiones | Araneae | Blattodea | Native Arthropods | Introduced Arthropods | Total Arthropods | Hemiptera | Rocks | Plants |
| 1 | 8.22.2020 | *Incilius alvarius* | NA | NA | NA | Adult | T | Native | Flushed | 7 | 0 | 0 | 0 | 0 | 0 | 0 | 0 | 7 | 0 | 7 | 0 | 1 | 0 |
| 2 | 8.22.2020 | *Anaxryus punctatus* | NA | NA | NA | Adult | T | Native | Flushed | 0 | 0 | 0 | 0 | 1 | 0 | 0 | 0 | 1 | 0 | 1 | 0 | 0 | 0 |
| 3 | 8.25.2020 | *Incilius alvarius* | 8.3 | NA | NA | Adult | A | Urban | Flushed | 3 | 0 | 0 | 0 | 15 | 0 | 0 | 2 | 18 | 2 | 20 | 0 | 0 | 1 |
| 4 | 8.27.2020 | *Incilius alvarius* | 13.6 | 265 | NA | Adult | A | Urban | Flushed | 0 | 1 | 0 | 0 | 0 | 0 | 0 | 0 | 1 | 0 | 1 | 0 | 0 | 0 |
| 5 | 8.27.2020 | *Incilius alvarius* | 10.4 | 115 | NA | Adult | A | Urban | Flushed | 0 | 0 | 0 | 0 | 0 | 0 | 0 | 0 | 0 | 0 | 0 | 0 | 0 | 0 |
| 6 | 8.27.2020 | *Incilius alvarius* | 14.8 | 400 | NA | Adult | A | Urban | Both | 0 | 0 | 1 | 0 | 0 | 0 | 0 | 2 | 1 | 2 | 3 | 0 | 1 | 1 |
| 7 | 8.27.2020 | *Incilius alvarius* | 13.9 | 360 | NA | Adult | A | Urban | Flushed | 1 | 1 | 0 | 0 | 0 | 0 | 0 | 0 | 2 | 0 | 2 | 0 | 0 | 1 |
| 8 | 8.27.2020 | *Incilius alvarius* | 12.8 | 230 | NA | Adult | A | Urban | Flushed | 0 | 1 | 0 | 0 | 0 | 0 | 0 | 0 | 1 | 0 | 1 | 0 | 0 | 0 |
| 9 | 8.27.2020 | *Incilius alvarius* | 2.5 | 0.5 | NA | Juvenile | A | Urban | Flushed | 0 | 0 | 0 | 0 | 0 | 0 | 0 | 0 | 0 | 0 | 0 | 0 | 0 | 0 |
| 10 | 8.27.2020 | *Anaxryus cognatus* | 8.9 | 88 | NA | Adult | A | Urban | Flushed | 0 | 1 | 0 | 0 | 0 | 0 | 0 | 0 | 1 | 0 | 1 | 0 | 1 | 1 |
| 11 | 8.25.2020 | *Anaxryus punctatus* | 5.4 | NA | NA | Adult | A | Urban | Flushed | 5 | 0 | 0 | 0 | 0 | 0 | 0 | 0 | 5 | 0 | 5 | 0 | 0 | 0 |
| 12 | 8.25.2020 | *Scaphiopus couchii* | 5.3 | NA | NA | Adult | A | Urban | Flushed | 0 | 0 | 0 | 0 | 0 | 0 | 0 | 0 | 0 | 0 | 0 | 0 | 0 | 0 |
| 13 | 8.28.2020 | *Scaphiopus couchii* | 5.1 | 23.5 | Female | Adult | M | Urban | Flushed | 4 | 0 | 2 | 0 | 7 | 0 | 0 | 0 | 14 | 0 | 14 | 1 | 1 | 0 |
| 14 | 8.28.2020 | *Scaphiopus couchii* | 5.1 | 38.5 | Male | Adult | M | Urban | Flushed | 2 | 0 | 0 | 0 | 1 | 0 | 0 | 1 | 4 | 0 | 4 | 0 | 0 | 0 |
| 15 | 8.28.2020 | *Scaphiopus couchii* | 4.8 | 20.5 | Male | Adult | M | Urban | Flushed | 2 | 0 | 0 | 2 | 0 | 0 | 0 | 0 | 2 | 2 | 4 | 0 | 0 | 0 |
| 16 | 8.28.2020 | *Incilius alvarius* | 9.7 | 125 | NA | Adult | M | Urban | Both | 6 | 0 | 1 | 0 | 3 | 0 | 1 | 0 | 11 | 0 | 11 | 0 | 0 | 2 |
| 17 | 8.28.2020 | *Anaxryus cognatus* | 7.1 | 41 | Male | Adult | M | Urban | Fecal | 0 | 0 | 0 | 0 | 5 | 0 | 0 | 0 | 5 | 0 | 5 | 0 | 0 | 0 |
| 18 | 8.28.2020 | *Anaxryus cognatus* | 9 | 75 | Female | Adult | M | Urban | Fecal | 0 | 0 | 0 | 0 | 5 | 0 | 0 | 0 | 5 | 0 | 5 | 0 | 0 | 0 |
| 19 | 8.29.2020 | *Incilius alvarius* | 15.2 | 360 | NA | Adult | M | Urban | Both | 3 | 0 | 0 | 0 | 14 | 0 | 0 | 0 | 17 | 0 | 17 | 0 | 0 | 4 |
| 20 | 8.29.2020 | *Incilius alvarius* | NA | NA | NA | Adult | M | Urban | Flushed | 2 | 0 | 0 | 0 | 2 | 0 | 0 | 0 | 4 | 0 | 4 | 0 | 0 | 0 |
| 21 | 8.30.2020 | *Anaxryus punctatus* | 6.1 | 20 | NA | Adult | M | Urban | Fecal | 0 | 0 | 0 | 0 | 1 | 0 | 0 | 0 | 1 | 0 | 1 | 0 | 0 | 2 |
| 22 | 8.30.2020 | *Anaxryus punctatus* | 5.5 | 14 | NA | Adult | M | Urban | Fecal | 4 | 0 | 0 | 0 | 2 | 0 | 0 | 0 | 6 | 0 | 6 | 0 | 0 | 6 |
| 23 | 8.30.2020 | *Anaxryus cognatus* | 8.4 | 83 | NA | Adult | M | Urban | Fecal | 1 | 0 | 0 | 0 | 12 | 0 | 0 | 0 | 13 | 0 | 13 | 0 | 0 | 1 |
| 24 | 8.30.2020 | *Anaxryus punctatus* | 4.8 | 14 | NA | Adult | M | Urban | Fecal | 0 | 0 | 0 | 0 | 3 | 0 | 0 | 0 | 3 | 0 | 3 | 0 | 0 | 0 |
| 25 | 8.30.2020 | *Incilius alvarius* | 4 | 6 | NA | Juvenile | M | Urban | Fecal | 6 | 0 | 0 | 0 | 0 | 0 | 0 | 0 | 6 | 0 | 6 | 0 | 0 | 2 |
| 26 | 8.30.2020 | *Incilius alvarius* | 10.5 | 105 | NA | Adult | M | Urban | Fecal | 3 | 0 | 0 | 0 | 0 | 0 | 0 | 0 | 3 | 0 | 3 | 0 | 0 | 1 |
| 27 | 8.30.2020 | *Incilius alvarius* | 6.8 | 27 | NA | Subadult | P | Native | Fecal | 1 | 0 | 0 | 0 | 0 | 0 | 0 | 0 | 1 | 0 | 1 | 0 | 0 | 0 |
| 28 | 8.30.2020 | *Incilius alvarius* | 8.2 | 65 | NA | Adult | M | Urban | Fecal | 1 | 0 | 0 | 0 | 0 | 0 | 0 | 0 | 1 | 0 | 1 | 0 | 2 | 6 |
| 29 | 8.30.2020 | *Scaphiopus couchii* | 5 | NA | Male | Adult | P | Native | Fecal | 1 | 0 | 0 | 0 | 0 | 0 | 0 | 0 | 1 | 0 | 1 | 0 | 0 | 1 |
| 30 | 8.30.2020 | *Scaphiopus couchii* | 6.3 | NA | Female | Adult | P | Native | Flushed | 0 | 0 | 0 | 0 | 0 | 0 | 0 | 0 | 0 | 0 | 0 | 0 | 0 | 0 |
| 31 | 8.30.2020 | *Scaphiopus couchii* | 5.6 | NA | Female | Adult | P | Native | Flushed | 0 | 0 | 0 | 0 | 0 | 0 | 0 | 0 | 0 | 0 | 0 | 0 | 0 | 0 |
| 32 | 8.30.2020 | *Scaphiopus couchii* | 3.8 | NA | Female | Subadult | P | Native | Flushed | 0 | 0 | 0 | 0 | 0 | 0 | 0 | 0 | 0 | 0 | 0 | 0 | 0 | 0 |
| 33 | 9.1.2020 | *Scaphiopus couchii* | 5.2 | 28 | Male | Adult | A | Urban | Flushed | 0 | 0 | 0 | 0 | 0 | 0 | 0 | 0 | 0 | 0 | 0 | 0 | 0 | 0 |
| 34 | 9.1.2020 | *Scaphiopus couchii* | 6 | 28 | Female | Adult | A | Urban | Flushed | 0 | 0 | 0 | 0 | 0 | 0 | 0 | 0 | 0 | 0 | 0 | 0 | 0 | 0 |
| 35 | 9.1.2020 | *Anaxryus punctatus* | 5.5 | 28 | NA | Adult | A | Urban | Both | 0 | 0 | 0 | 0 | 0 | 0 | 1 | 0 | 1 | 0 | 1 | 0 | 0 | 0 |
| 36 | 9.1.2020 | *Incilius alvarius* | 5.1 | 12 | NA | Subadult | A | Urban | Fecal | 0 | 0 | 0 | 0 | 0 | 0 | 0 | 0 | 0 | 0 | 0 | 0 | 0 | 8 |
| 37 | 9.1.2020 | *Incilius alvarius* | 3.4 | 3 | NA | Juvenile | A | Urban | Flushed | 0 | 0 | 0 | 0 | 2 | 0 | 0 | 1 | 2 | 1 | 3 | 0 | 1 | 0 |
| 38 | 9.1.2020 | *Incilius alvarius* | 3.4 | <5 | NA | Juvenile | A | Urban | Fecal | 0 | 0 | 0 | 0 | 10 | 0 | 0 | 0 | 10 | 0 | 10 | 0 | 0 | 2 |
| 39 | 9.1.2020 | *Incilius alvarius* | 2.8 | <5 | NA | Juvenile | A | Urban | Flushed | 0 | 0 | 0 | 0 | 0 | 0 | 0 | 0 | 0 | 0 | 0 | 0 | 0 | 0 |
| 40 | 9.3.2020 | *Incilius alvarius* | 13.4 | 310 | NA | Adult | T | Native | Flushed | 0 | 0 | 0 | 0 | 1 | 0 | 0 | 0 | 1 | 0 | 1 | 0 | 0 | 0 |
| 41 | 9.3.2020 | *Incilius alvarius* | 14.3 | 320 | NA | Adult | T | Native | Both | 12 | 0 | 0 | 0 | 0 | 0 | 0 | 0 | 12 | 0 | 12 | 0 | 0 | 5 |
| 42 | 9.3.2020 | *Incilius alvarius* | 11.2 | 175 | NA | Adult | T | Native | Flushed | 3 | 0 | 0 | 0 | 0 | 0 | 0 | 0 | 3 | 0 | 3 | 0 | 15 | 4 |
| 43 | 9.3.2020 | *Incilius alvarius* | 16.2 | 392 | NA | Adult | T | Native | Both | 4 | 0 | 0 | 0 | 0 | 0 | 0 | 0 | 4 | 0 | 4 | 0 | 1 | 1 |
| 44 | 9.3.2020 | *Incilius alvarius* | 13.9 | 295 | NA | Adult | T | Native | Both | 2 | 0 | 0 | 0 | 80 | 0 | 1 | 0 | 83 | 0 | 83 | 0 | 0 | 2 |
| 45 | 9.3.2020 | *Incilius alvarius* | 11.4 | 210 | NA | Adult | T | Native | Both | 1 | 0 | 0 | 0 | 8 | 1 | 0 | 0 | 10 | 0 | 10 | 0 | 1 | 0 |
| 46 | 9.6.2020 | *Incilius alvarius* | NA | NA | NA | Adult | T | Native | Flushed | 8 | 0 | 0 | 0 | 0 | 0 | 0 | 0 | 8 | 0 | 8 | 0 | 0 | 0 |
| 47 | 9.6.2020 | *Anaxryus punctatus* | 5 | 5.6 | Male | Adult | T | Native | Flushed | 0 | 0 | 0 | 0 | 0 | 0 | 0 | 0 | 0 | 0 | 0 | 0 | 0 | 0 |
| 49 | 9.6.2020 | *Incilius alvarius* | NA | NA | NA | Adult | T | Native | Flushed | 0 | 0 | 0 | 0 | 0 | 0 | 0 | 0 | 0 | 0 | 0 | 0 | 0 | 0 |
| 50 | 9.10.2020 | *Incilius alvarius* | 13.2 | 240 | NA | Adult | G | Native | Fecal | 5 | 0 | 0 | 0 | 0 | 0 | 0 | 0 | 5 | 0 | 5 | 0 | 0 | 0 |
