## Supplemental Table 4 for "Diet and chemical defenses of the Sonoran Desert toads"

| Anuran | | Sample Identifier (Anuran Number_Sample, FE = fecal) | Habitat | Identification Method | Sanger Results | | Consensus Identification | | | | | Additional Notes |
|---|---|---|---|---|---|---|---|---|---|---|---|---|
| Identification Number | Species | | | | BLASTn match GenBank Accession (if applicable) | Similarity (if applicable), threshold = 98% | Family | Genus | Genus Species | Final Prey Identification | Identified to | |
| 1 | *Incilius alvarius* | 1_a | Native | Visual | NA | NA | Elateridae | *Melanotus* | *Melanotus cribricollis* | *Melanotus cribricollis* | Species | |
| 1 | *Incilius alvarius* | 1_b | Native | Both | HM433098 | 98.406 | Tenebrionidae | | | Alleculinae | Subfamily | |
| 1 | *Incilius alvarius* | 1_c | Native | Both | HM433098 | 98.381 | Tenebrionidae | | | Alleculinae | Subfamily | |
| 1 | *Incilius alvarius* | 1_d | Native | Both | HM433098 | 98.333 | Tenebrionidae | | | Alleculinae | Subfamily | |
| 1 | *Incilius alvarius* | 1_e | Native | Visual | HM433098 | 97.927 | Tenebrionidae | | | Alleculinae | Subfamily | |
| 1 | *Incilius alvarius* | 1_g | Native | Visual | HM433098 | 99.346 | Tenebrionidae | *Hymenorus* | | *Hymenorus sp.* | Genus | Sanger matched to subfamily (Alleculinae) only |
| 2 | *Anaxyrus punctatus* | 2_a | Native | Visual | NA | NA | Formicidae | | | Formicidae | Family | Sanger yielded Pheidole obtusopinosa at 97% (100% reverese strand) |
| 3 | *Incilius alvarius* | 03_a | Urban | Sanger | MT465995 | 100 | Blattidae | *Shelfordella* | *Shelfordella lateralis* | *Shelfordella lateralis* | Species | |
| 3 | *Incilius alvarius* | 03_c | Urban | Visual | NA | NA | Formicidae | *Pogonomyrmex* | *Pogonomyrmex cf rugosus* | *Pogonomyrmex cf rugosus* | Species | |
| 3 | *Incilius alvarius* | 03_d | Urban | Visual | NA | NA | Salpingidae | *Dacoderus* | | *Dacoderus sp.* | Genus | |
| 3 | *Incilius alvarius* | 03_e | Urban | Both | HQ984105 | 99.127 | Chrysomelidae | | | Bruchinae | Subfamily | |
| 3 | *Incilius alvarius* | 03_g | Urban | Sanger | MF149455 | 100 | Blattidae | *Periplaneta* | *Periplaneta americana* | *Periplaneta americana* | Species | |
| 3 | *Incilius alvarius* | 03_h | Urban | Both | HQ984105 | 99.593 | Chrysomelidae | | | Bruchinae | Subfamily | |
| 6 | *Incilius alvarius* | 06_d | Urban | Sanger | MZ440657 | 100 | Gryllidae | *Gryllodes* | | *Gryllodes sp.* | Genus | |
| 6 | *Incilius alvarius* | 06_f_FE_mix | Urban | Both | HQ984105 | 99.593 | Blattidae | | | Blattidae | Family | |
| 7 | *Incilius alvarius* | 07_a | Urban | Sanger | MG230727 | 99.16 | Elateridae | *Horistonotus* | *Horistonotus cf. simplex* | *Horistonotus cf. simplex* | Species | Visually identified as Horistonotus simplex or Esthesopus parcus |
| 10 | *Anaxyrus cognatus* | 10_a | Urban | Sanger | OP628433 | 100 | Libellulidae | *Pantala* | *Pantala flavescens* | *Pantala flavescens* | Species | |
| 11 | *Anaxyrus punctatus* | 11_a | Urban | Visual | NA | NA | Hydrophilidae | *Enochrus* | | *Enochrus sp.* | Genus | This and following Enochrus sp. identified as Enochrus (Lumetus) falcarius by Sanger Sequencing, but with low similarity (<91%) |
| 11 | *Anaxyrus punctatus* | 11_b | Urban | Visual | NA | NA | Hydrophilidae | *Enochrus* | | *Enochrus sp.* | Genus | |
| 11 | *Anaxyrus punctatus* | 11_c | Urban | Visual | NA | NA | Hydrophilidae | *Enochrus* | | *Enochrus sp.* | Genus | |
| 11 | *Anaxyrus punctatus* | 11_d | Urban | Visual | NA | NA | Hydrophilidae | *Enochrus* | | *Enochrus sp.* | Genus | |
| 11 | *Anaxyrus punctatus* | 11_e | Urban | Visual | NA | NA | Hydrophilidae | *Enochrus* | | *Enochrus sp.* | Genus | |
| 13 | *Scaphiopus couchii* | 13_a | Urban | Both | HQ928784 | 98.696 | Formicidae | *Camponotus* | | *Camponotus sp.* | Genus | Visually identified as member of festinatus species complex |
| 13 | *Scaphiopus couchii* | 13_b | Urban | Visual | NA | NA | Cicadellidae | *Dixianus* | *Dixianus utahnus* | *Dixianus utahnus* | Species | |
| 13 | *Scaphiopus couchii* | 13_c | Urban | Sanger | MW340162 | 100 | Carabidae | *Discoderus* | *Discoderus robustus* | *Discoderus robustus* | Species | Visually identified as Discoderus sp. |
| 13 | *Scaphiopus couchii* | 13_f | Urban | Both | KT006489 | 100 | Formicidae | *Pogonomyrmex* | *Pogonomyrmex cf rugosus* | *Pogonomyrmex cf rugosus* | Species | |
| 13 | *Scaphiopus couchii* | 13_g | Urban | Both | KT006489 | 100 | Formicidae | *Pogonomyrmex* | *Pogonomyrmex cf rugosus* | *Pogonomyrmex cf rugosus* | Species | |
| 13 | *Scaphiopus couchii* | 13_h | Urban | Both | KT006489 | 100 | Formicidae | *Pogonomyrmex* | *Pogonomyrmex cf rugosus* | *Pogonomyrmex cf rugosus* | Species | |
| 13 | *Scaphiopus couchii* | 13_i | Urban | Both | KT006489 | 100 | Formicidae | *Pogonomyrmex* | *Pogonomyrmex cf rugosus* | *Pogonomyrmex cf rugosus* | Species | |
| 13 | *Scaphiopus couchii* | 13_j | Urban | Sanger | AF317179 | 100 | Acrididae | *Melanoplus* | *Melanoplus aridus* | *Melanoplus aridus* | Species | |
| 13 | *Scaphiopus couchii* | 13_l | Urban | Visual | NA | NA | Scarabaeidae | | | Aphodiinae | Subfamily | |
| 14 | *Scaphiopus couchii* | 14_a | Urban | Visual | NA | NA | Elateridae | *Aeolus* | | *Aeolus sp.* | Genus | |
| 14 | *Scaphiopus couchii* | 14_c | Urban | Visual | NA | NA | Formicidae | *Pogonomyrmex* | *Pogonomyrmex cf rugosus* | *Pogonomyrmex cf rugosus* | Species | |
| 14 | *Scaphiopus couchii* | 14_d | Urban | Visual | NA | NA | | | | Isoptera | Infraorder | |
| 15 | *Scaphiopus couchii* | 15_a | Urban | Visual | NA | NA | Carabidae | | | Carabidae | Family | Visually identified as Discoderus sp. (tentative) |
| 15 | *Scaphiopus couchii* | 15_b | Urban | Both | KC014537 | 100 | Oligotomidae | *Oligotoma* | *Oligotoma nigra* | *Oligotoma nigra* | Species | |
| 15 | *Scaphiopus couchii* | 15_c | Urban | Visual | NA | NA | Oligotomidae | *Oligotoma* | *Oligotoma nigra* | *Oligotoma nigra* | Species | |
| 16 | *Incilius alvarius* | 16_a | Urban | Visual | NA | NA | Philodromidae | *Tibellus* | | *Tibellus sp.* | Genus | |
| 16 | *Incilius alvarius* | 16_b | Urban | Sanger | HQ983698 | 99.597 | Acrididae | *Psoloessa* | *Psoloessa texana* | *Psoloessa texana* | Species | |
| 16 | *Incilius alvarius* | 16_l_FE | Urban | Both | HM386206 | 97.38 | Tenebrionidae | *Eleodes* | | *Eleodes sp.* | Genus | Sanger identification was E. carbonaria |
| 19 | *Incilius alvarius* | 19_a | Urban | Visual | NA | NA | Formicidae | *Pogonomyrmex* | *Pogonomyrmex cf rugosus* | *Pogonomyrmex cf rugosus* | Species | |
| 19 | *Incilius alvarius* | 19_u_FE | Urban | Visual | NA | NA | Tenebrionidae | *Eleodes* | | *Eleodes sp.* | Genus | |
| 19 | *Incilius alvarius* | 19_v_FE | Urban | Visual | NA | NA | Tenebrionidae | *Conibius* | | *Conibius sp.* | Genus | |
| 19 | *Incilius alvarius* | 19_w_FE | Urban | Visual | NA | NA | Tenebrionidae | *Eleodes* | | *Eleodes sp.* | Genus | |
| 20 | *Incilius alvarius* | 20_a | Urban | Visual | NA | NA | Formicidae | *Pogonomyrmex* | *Pogonomyrmex cf rugosus* | *Pogonomyrmex cf rugosus* | Species | |
| 20 | *Incilius alvarius* | 20_b | Urban | Visual | NA | NA | Formicidae | *Pogonomyrmex* | *Pogonomyrmex cf rugosus* | *Pogonomyrmex cf rugosus* | Species | Identified as P. rugosus from Sanger Sequencing, but low similarity (93%) |
| 23 | *Anaxyrus cognatus* | 23_FE_a | Urban | Visual | NA | NA | Formicidae | *Pogonomyrmex* | *Pogonomyrmex cf rugosus* | *Pogonomyrmex cf rugosus* | Species | |
| 26 | *Incilius alvarius* | 26_a_FE | Urban | Visual | NA | NA | Tenebrionidae | *Eleodes* | | *Eleodes sp.* | Genus | |
| 26 | *Incilius alvarius* | 26_c_FE | Urban | Visual | NA | NA | Tenebrionidae | *Eleodes* | | *Eleodes sp.* | Genus | |
| 26 | *Incilius alvarius* | 26_d_FE | Urban | Visual | NA | NA | Tenebrionidae | *Eleodes* | | *Eleodes sp.* | Genus | |
| 40 | *Incilius alvarius* | 40_a_FE | Native | Visual | NA | NA | Vespidae | | | Vespidae | Family | |
| 40-45 | *Incilius alvarius* | 40-45_g_FE | Native | Visual | NA | NA | Tenebrionidae | | | Tenebrionidae | Family | Fecal samples of uncertain origin |
| 40-46 | *Incilius alvarius* | 40-45_o_FE | Native | Visual | NA | NA | Tenebrionidae | *Eleodes* | | *Eleodes sp.* | Genus | Fecal samples of uncertain origin |
| 41 | *Incilius alvarius* | 41_d | Native | Visual | NA | NA | Tenebrionidae | | | Alleculinae | Subfamily | |
| 41 | *Incilius alvarius* | 41_e | Native | Both | HM433098 | 98.298 | Tenebrionidae | | | Alleculinae | Subfamily | |
| 41 | *Incilius alvarius* | 41_f | Native | Both | HM433098 | 98.326 | Tenebrionidae | | | Alleculinae | Subfamily | |
| 41 | *Incilius alvarius* | 41_g | Native | Both | HM433098 | 98.381 | Tenebrionidae | | | Alleculinae | Subfamily | |
| 41 | *Incilius alvarius* | 41_h | Native | Visual | HM433098 | 97.951 | Tenebrionidae | | | Alleculinae | Subfamily | Sanger identification as Alleculinae sp. just below 98% threshold |
| 42 | *Incilius alvarius* | 42_b_mix | Native | Visual | NA | NA | Tenebrionidae | | | Tenebrionidae | Family | |
| 43 | *Incilius alvarius* | 43_d_FE | Native | Visual | NA | NA | Tenebrionidae | *Cryptoglossa* | *Cryptoglossa variolosa* | *Cryptoglossa variolosa* | Species | |
| 43 | *Incilius alvarius* | 43_e_FE | Native | Visual | NA | NA | Tenebrionidae | *Cryptoglossa* | *Cryptoglossa variolosa* | *Cryptoglossa variolosa* | Species | |
| 43 | *Incilius alvarius* | 43_g_FE | Native | Visual | NA | NA | Tenebrionidae | *Eleodes* | | *Eleodes sp.* | Genus | |
| 43 | *Incilius alvarius* | 43_i_FE | Native | Visual | NA | NA | Tenebrionidae | *Cryptoglossa* | *Cryptoglossa variolosa* | *Cryptoglossa variolosa* | Species | |
| 44 | *Incilius alvarius* | 44_d | Native | Visual | NA | NA | Corinnidae | *Septentrinna* | | *Septentrinna sp.* | Genus | |
| 44 | *Incilius alvarius* | 44_e | Native | Both | KJ141939 | 100 | Formicidae | *Pheidole* | *Pheidole rhea* | *Pheidole rhea* | Species | |
| 44 | *Incilius alvarius* | 44_l | Native | Visual | NA | NA | Formicidae | *Pheidole* | *Pheidole rhea* | *Pheidole rhea* | Species | |
| 44 | *Incilius alvarius* | 44_m | Native | Both | KJ141941 | 100 | Formicidae | *Pheidole* | *Pheidole rhea* | *Pheidole rhea* | Species | |
| 44 | *Incilius alvarius* | 44_n | Native | Visual | NA | NA | Formicidae | *Pheidole* | *Pheidole rhea* | *Pheidole rhea* | Species | |
| 44 | *Incilius alvarius* | 44_o | Native | Visual | NA | NA | Formicidae | *Pheidole* | *Pheidole rhea* | *Pheidole rhea* | Species | |
| 44 | *Incilius alvarius* | 44_p | Native | Visual | NA | NA | Formicidae | *Pheidole* | *Pheidole rhea* | *Pheidole rhea* | Species | |
| 44 | *Incilius alvarius* | 44_q | Native | Visual | NA | NA | Formicidae | *Pheidole* | *Pheidole rhea* | *Pheidole rhea* | Species | |
| 44 | *Incilius alvarius* | 44_r | Native | Visual | NA | NA | Formicidae | *Pheidole* | *Pheidole rhea* | *Pheidole rhea* | Species | |
| 44 | *Incilius alvarius* | 44_z_FE | Native | Visual | NA | NA | Tenebrionidae | *Argoporis* | | *Argoporis sp.* | Genus | |
| 45 | *Incilius alvarius* | 45_a | Native | Both | AY649857 | 99.62 | Buthidae | *Centruroides* | *Centruroides sculpturatus* | *Centruroides sculpturatus* | Species | |
| 45 | *Incilius alvarius* | 45_c | Native | Both | KJ141940 | 100 | Formicidae | *Pheidole* | *Pheidole rhea* | *Pheidole rhea* | Species | |
| 45 | *Incilius alvarius* | 45_c_2 | Native | Visual | NA | NA | Formicidae | *Solenopsis* | | *Solenopsis sp.* | Genus | From same mixed sample as 45_c |
| 46 | *Incilius alvarius* | 46_a | Native | Both | HM433098 | 99.624 | Tenebrionidae | | | Alleculinae | Subfamily | |
| 46 | *Incilius alvarius* | 46_b | Native | Both | HM433098 | 98.406 | Tenebrionidae | | | Alleculinae | Subfamily | |
| 46 | *Incilius alvarius* | 46_c | Native | Visual | HM433098 | 97.619 | Tenebrionidae | | | Alleculinae | Subfamily | |
| 46 | *Incilius alvarius* | 46_d | Native | Both | HM433098 | 98.361 | Tenebrionidae | | | Alleculinae | Subfamily | |
| 46 | *Incilius alvarius* | 46_e | Native | Both | HM433098 | 98.374 | Tenebrionidae | | | Alleculinae | Subfamily | |
| 46 | *Incilius alvarius* | 46_f | Native | Both | HM433098 | 98.406 | Tenebrionidae | | | Alleculinae | Subfamily | |
| 50 | *Incilius alvarius* | 50_a_FE | Native | Visual | NA | NA | Tenebrionidae | | | Tenebrionidae | Family | |
| 50 | *Incilius alvarius* | 50_b_FE | Native | Visual | NA | NA | Tenebrionidae | | | Tenebrionidae | Family | |
| 50 | *Incilius alvarius* | 50_c_FE | Native | Visual | NA | NA | Tenebrionidae | | | Tenebrionidae | Family | |
| 50 | *Incilius alvarius* | 50_d_FE | Native | Visual | NA | NA | Tenebrionidae | | | Tenebrionidae | Family | |
| 50 | *Incilius alvarius* | 50_e_FE | Native | Visual | NA | NA | Tenebrionidae | | | Tenebrionidae | Family | |
| 50 | *Incilius alvarius* | 50_f_FE | Native | Visual | NA | NA | Tenebrionidae | | | Tenebrionidae | Family | |
